# Identification and evolution of avian endogenous foamy viruses

**DOI:** 10.1101/707083

**Authors:** Yicong Chen, Xiaoman Wei, Guojie Zhang, Edward C. Holmes, Jie Cui

## Abstract

A history of long-term co-divergence means that foamy viruses (family *Retroviridae*) provide an ideal framework to understanding virus-host evolution over extended time periods. Endogenous foamy viruses (EndFVs) are rare, and to date have only been described in a limited number of mammals, amphibians, reptiles and fish genomes. By screening 414 avian genomes we identified EndFVs in two bird species: the Maguari Stork (*Ciconia maguari*) and the Oriental Stork (*C. boyciana*). Analyses of phylogenetic relationships, genome structures and flanking sequences revealed a single origin of EndFVs in *Ciconia* species. In addition, the marked incongruence between the virus and host phylogenies suggested that this integration event occurred independently in birds. In sum, by providing evidence that birds can be infected with foamy viruses, we fill the last major gap in the taxonomic distribution of foamy viruses and their animal hosts.

## Introduction

Retroviruses (family *Retroviridae*) are viruses of substantial medical and economic significance as some are associated with severe infectious disease or are oncogenic (Hayward, et al. 2015; Aiewsakun and Katzourakis 2017; Xu, et al. 2018). Retroviruses are also of evolutionary importance as they have occasionally invaded the host germ line, leading to the generation of endogenous retroviruses (ERVs) and hence genomic ‘fossils’ (Stoye 2012; Johnson 2015, 2019). ERVs are widely distributed in vertebrates (Hayward, et al. 2013; Cui, et al. 2014; Hayward, et al. 2015; Xu, et al. 2018) and provide important insights into the origin and long-term evolution of retroviruses. However, some complex retroviruses such as lenti-, delta- and spuma viruses, only relatively rarely appear as endogenized forms.

As well leaving a litany of endogenous copies in host genomes, foamy viruses are of particular importance because they exhibit a history of long-term co-divergence with their vertebrate hosts (Switzer, et al. 2005). Endogenous foamy viruses (EndFVs) were first discovered in sloths (Katzourakis, et al. 2009), and then found in several primate genomes (Han and Worobey 2012b, 2014; Katzourakis, et al. 2014). The subsequent discovery of EndFV and EndFV-like copies in fish genomes indicated that foamy viruses may have a deep evolutionary history within the vertebrates (Han and Worobey 2012a; Ruboyianes and Worobey 2016; Aiewsakun and Katzourakis 2017). Recently, three novel endogenous foamy viruses were identified in reptile genomes, although there is disagreement over their origins with some suggesting virus-host co-divergence over many millions of years (Wei et al. 2019), and others favoring cross-species transmission events (Aiewsakun, et al. 2019; Wei, et al. 2019). To date, no EndFVs have been identified in avian genomes.

## Materials and Methods

### Genome screening and viral genome structure identification

All 147 avian genomes available in GenBank as of June 2019 (Supplementary Table S1) and 267 genomes from the ‘Bird 10K’ program were screened for endogenous foamy viruses using the TBLASTN algorithm (Altschul, et al. 1990) and the protein sequences of representative exogenous foamy viruses, EndFVs and endogenous foamy-like viruses (Supplementary Table S2). A 35% sequence identity over a 30% region with an e-value set to 0.0001 was used to filter significant hits (Supplementary Table S3). Viral hits within large scaffolds (>20 kb) were assumed to represent *bona fide* ERVs. We then extended the flanking sequence of these hits to identify the viral long terminal repeats (LTRs) using BLASTN (Altschul, et al. 1990), LTR Finder (Xu and Wang 2007) and LTRharvest (Ellinghaus, et al. 2008). In accordance with the nomenclature proposed for ERVs (Gifford, et al. 2018), EndFVs were identified in the genomes of the Maguari Stork (*Ciconia maguari*) and the Oriental stork (*C. boyciana*). These were termed ‘ERV-Spuma.n-Cma’ and ‘ERV-Spuma.n-Cbo’, respectively (in which n represents the number of the viral sequences extracted from host genome) (Supplementary Table S4). Putative genome structures and conserved EndFV domains were identified using BLASTP, CD-search (Marchler-Bauer and Bryant 2004; Marchler-Bauer, et al. 2017) and ORFfinder (https://www.ncbi.nlm.nih.gov/orffinder/) at NCBI.

### Molecular dating

ERV integration time can be approximately estimated using the relation T = (D/R)/2, in which T is the integration time (million years, MY), D is the number of nucleotide differences per site between the pairwise LTRs, and R is the genomic substitution rate (nucleotide substitutions per site, per year). We used the previously estimated neutral nucleotide substitution rate for birds (1. 9 × 10^−9^ nucleotide substitutions per site, per year) (Zhang, et al. 2014). Two full-length ERVs-Spuma-Cma containing a pairwise intact LTRs were used to estimate integration time in this manner (Supplementary Table S5). We excluded ERV-Spum.1-Cbo from this dating exercise due to its defective 5’ LTR.

### Phylogenetic analysis

To describe the evolutionary relationship of EndFVs to other representative retroviruses, sequences of the Pol (Supplementary data set S1) and concatenated Gag-Pol-Env proteins (Supplementary data set S2) were aligned using MAFFT 7.222 (Katoh and Standley 2013) and confirmed manually in MEGA X (Kumar, et al. 2018). A phylogenetic tree of these data was then inferred using the maximum likelihood (ML) method in PhyML 3.1 (Guindon, et al. 2010), incorporating 100 bootstrap replicates to assess node robustness. The best-fit LG+Γ+I+F of amino acid substitution was selected for both Pol and concatenated Gag-Pol-Env protein data sets using ProtTest (Abascal, et al. 2005).

### Recombination analysis

To test for recombination in these data, we: (i) compared target site duplications (TSD) flanking the ERVs, as it has previously been shown that ERVs not flanked by the same TSDs likely arose by provirus recombination (Hughes and Coffin 2001), and (ii) screened for recombination in the pol and gag-pol-env nucleotide sequences of mammalian, tuatara, amphibian, lobe-finned fish and avian FVs/EndFVs using the Recombination Detection Program 4 (RDP4) (Martin, et al. 2015)

## Results and Discussion

### Discovery and characterization of endogenous foamy viral elements in avian genomes

To identify potential foamy (-like) viral elements in birds, we collected 147 available bird genomes from GenBank (Supplementary Table 1) and 267 genomes from the ‘Bird 10K’ project (Zhang, et al. 2015) and performed *in silico* TBLASTN, using the amino acid sequences of representative retroviruses as queries (Supplementary Table 2). This genomic mining identified 16 significant hits in the Maguari Stork and the 12 in Oriental Stork (Supplementary Table 3). We designated these ERV-Spuma.n-Cma and Spuma.n-Cbo, respectively (Gifford, et al. 2018) (Supplementary Table S3, Table S4). We considered hits within large scaffolds (>20 kilobases in length) to represent *bona fide* ERVs. We then extended the flanking sequences of these EndFVs on both sides to search for LTRs, as these define the boundary of the viral elements. Through this analysis we discovered two full-length EndFVs in the Maguari stork genome and one in the Oriental stork genome. The low copy number of EndFVs found in both two bird species accords with the observation that avian genomes generally harbor small numbers of endogenous viruses (Cui, et al. 2014). To further elucidate the relationship between these novel avian EndFVs and other retroviruses, Pol gene sequences (>500 amino acid residue in length) were used in a phylogenetic analysis (Fig. 1). Accordingly, our maximum likelihood phylogenetic tree revealed that the EndFVs discovered in birds formed a close and well supported monophyletic group within the foamy virus clade compatible with the idea that these avian EndFVs might have a single origin. Notably, however, because they were most closely related to the endogenous foamy viruses found in mammals rather than to those found in reptiles, the phylogenetic position of the avian EndFVs described here was incongruent with that of the host phylogeny (although the node associated with the tuatara EndFV was relatively poorly supported) (Fig. 2). This, and the overall rarity of EndFVs in birds, suggests that these avian EndFVs have an independent origin in birds and were not acquired through virus-host co-divergence, such that a non-avian retrovirus jumped into birds at some point during evolutionary history. No evidence for recombination was found in these data.

**Fig. 1.**
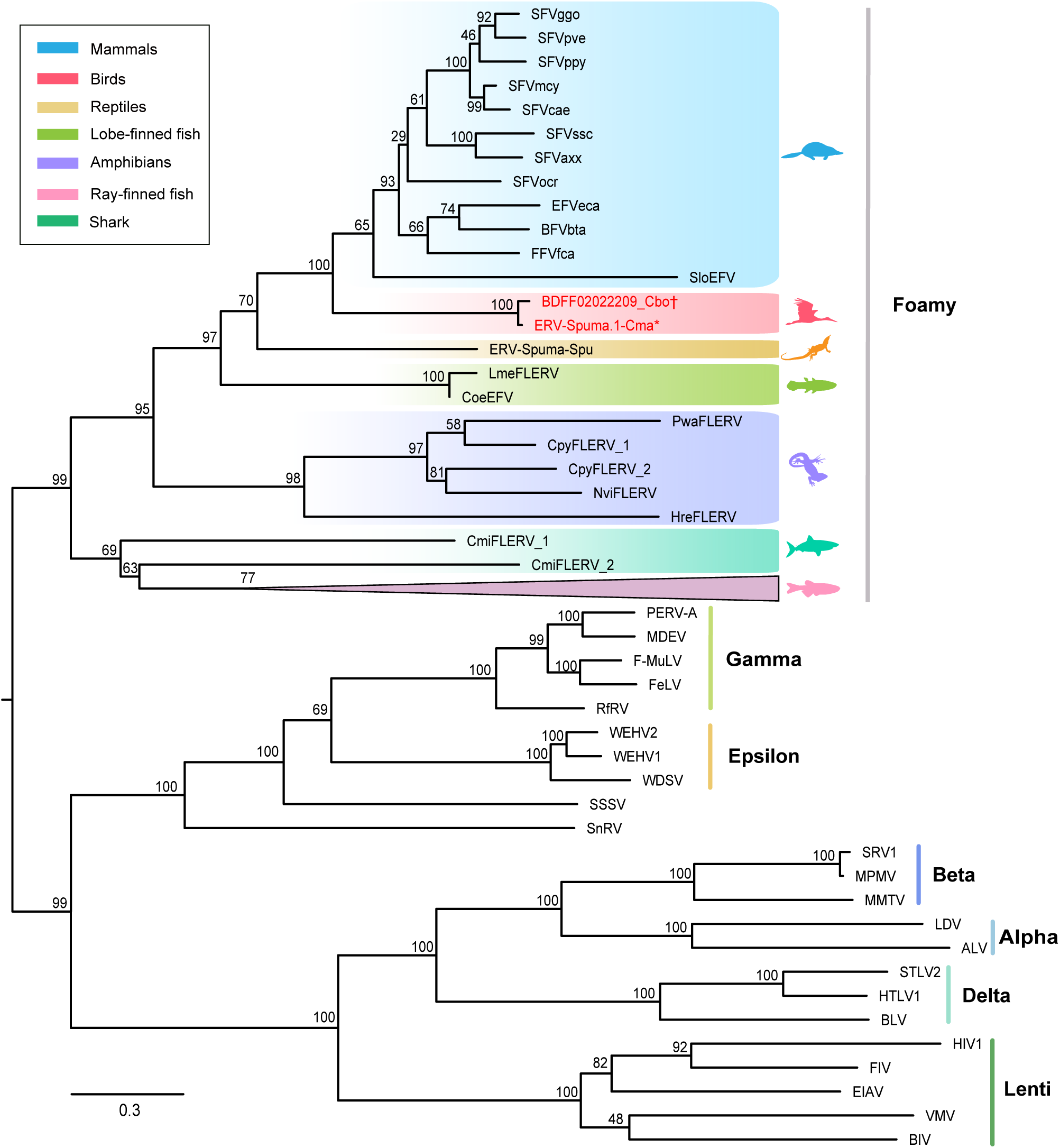
Phylogenetic tree of retroviruses and endogenous retroviruses, including the endogenous foamy viruses found in avian genomes. The tree was inferred using amino acid sequences of the Pol gene, and rooted using the remaining retroviral taxa (excluding the spumaviruses). The newly identified viral elements are labelled in red. * indicates the EndFV found in the Maguari Stork genome, while † denotes the EndFV from the Oriental Stork genome. The scale bar indicates the number of amino acid changes per site.

**Fig. 2.**
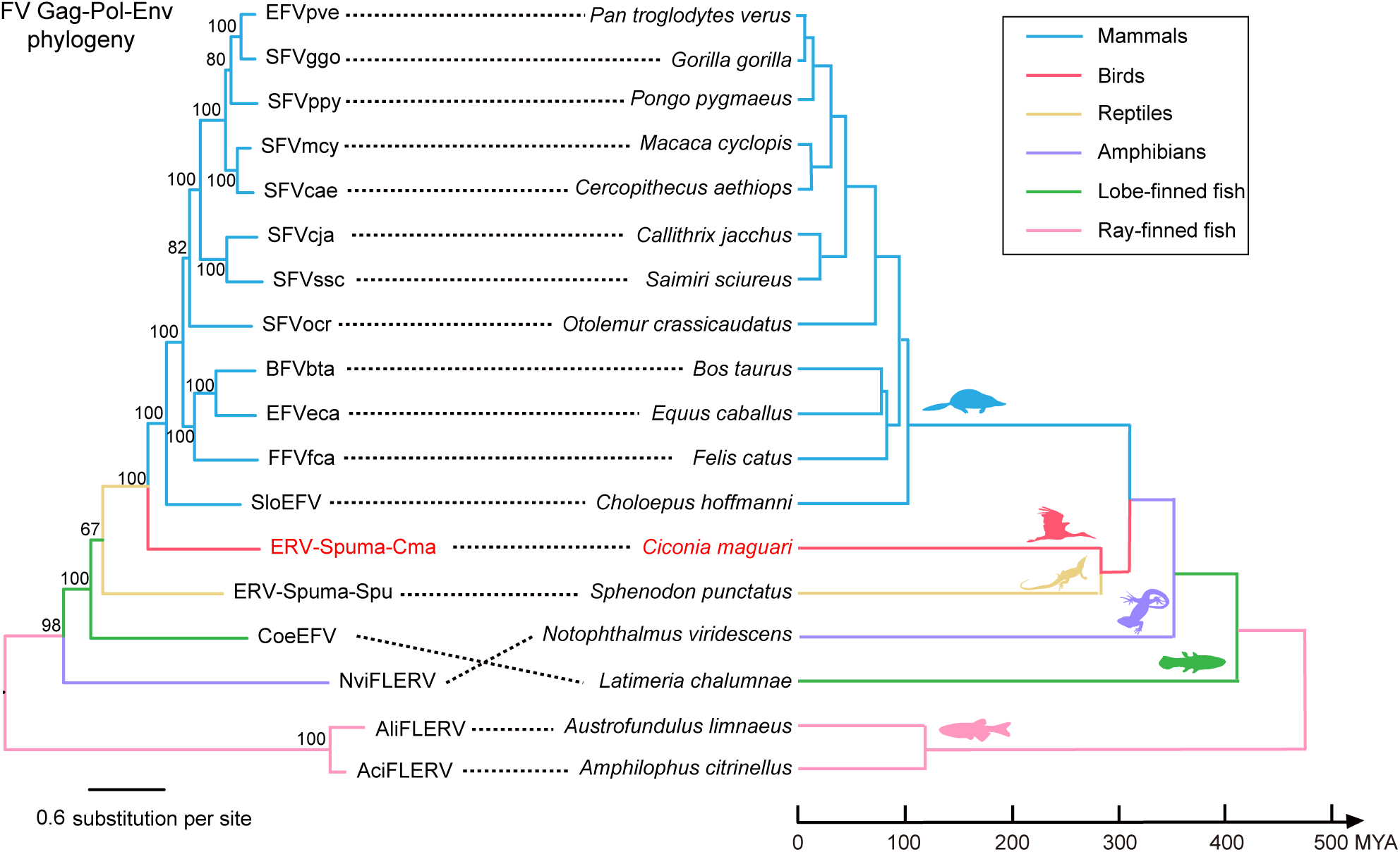
Evolutionary history of foamy viruses (left) and their vertebrate hosts (right). Associations between foamy viruses and their hosts are indicated by connecting lines. The avian EndFV and its host are labelled in red. Note that the avian EndFVs are more closely related to mammalian foamy viruses than the reptilian EndFV. Scale bars indicate the number of amino acid changes per site in the viruses and the host divergence times (million years ago, MYA).

### Genomic structure characterization

By searching for conserved domains against the Conserved Domains Database (www.ncbi.nlm.nih.gov/Structure/cdd), we identified three typical foamy conserved domains in the three full-length avian EndFVs: (1) the Spuma virus Gag domain (pfam03276) (Winkler, et al. 1997), (2) the Spuma aspartic protease (A9) domain (pfam03539) that is present in all mammalian foamy virus Pol proteins (Aiewsakun and Katzourakis 2017), and the (3) foamy virus envelope protein domain (pfam03408) (Han and Worobey 2012a; Wei, et al. 2019) (Supplementary Fig. S2). Furthermore, we identified an open reading frame 1 (ORF1) as an accessary gene in all three full-length avian EndFV genomes. These ORF1 sequences exhibited high nucleotide similarity (>88%) with each other, although not to any known accessory genes from foamy viruses (and the ORF1 in ERV-Spuma.1-Cma was split in two, due to two insertions and a nonsense mutation; Supplementary Fig. S1). Interestingly, the internal promoter of this ORF was located between 3’ end of the *env* gene and the 5’ start of the ORF gene. This is inconsistent with exogenous foamy viruses whose internal promoters are located toward the 3’ end of the *env* gene (Campbell, et al. 1994; Lochelt, et al. 153 1995).

In summary, the genomes of the new EndFVs documented here contained a pair of LTRs, although with no sequence similarity to other EndFV LTRs, and exhibited a typical spuma virus structure, with three main open reading frames (ORF) - gag, pol, and env - and one putative additional accessory gene, ORF 1 (Fig. 3, Supplementary Fig. S1).

**Fig. 3.**
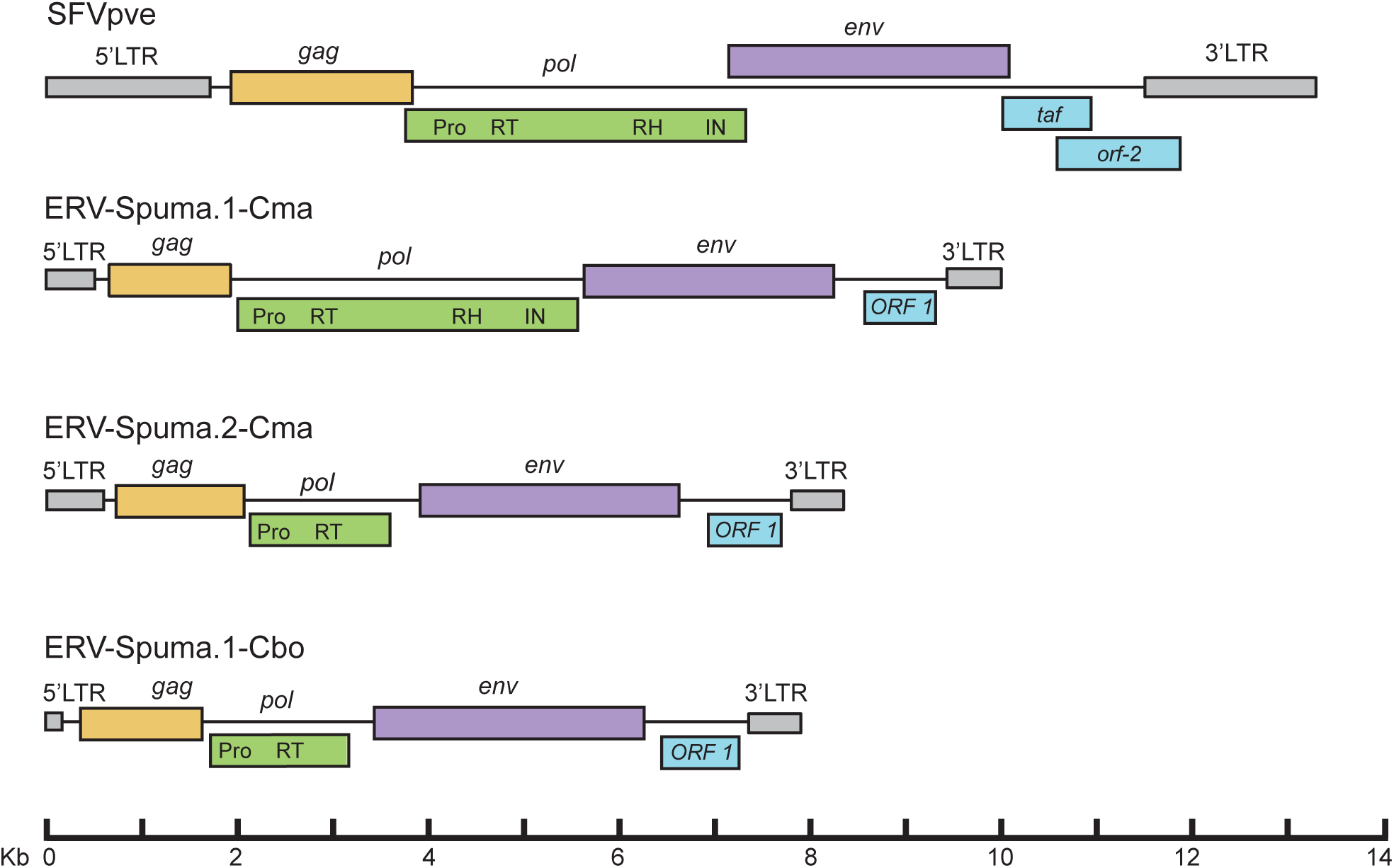
Genomic organization of exogenous foamy viruses (acc: NC_001364) and avian endogenous foamy viruses. LTR, long terminal repeat; Pro, protease; RT, reverse transcriptase; RH, ribonuclease H; IN, integrase.

### Vertical transmission of bird EndFVs

Surprisingly, ERV-Spuma.2-Cma and ERV-Spuma.1-Cbo shared 98% nucleotide sequence identity and contained the same deletion in the pol gene (Supplementary Fig. S3). By comparing the flanking sequences of these two EndFVs using BLASTN, we discovered that two scaffolds containing EndFV (BDFF02011124.1 in the Oriental stork and scaffold3222 in the Maguari Stork) shared 99% coverage with 98.66% sequence identity (e-value = 0.0). Furthermore, upon the ERV insertion, the target DNA fragment is also duplicated, resulting in target site duplication (TSDs) that differs among ERV insertions (Hughes and Coffin 2001). We were able to identify the same TSD flanking these two avian EndFVs (Supplementary Fig. S3), confirming that they have been vertically inherited. As such, we can infer that the most recent common ancestor of Maguari Stork and Oriental Stork, estimated to have existed between 3.2-13.2 million years ago (Jetz, et al. 2012), carried these EndFVs. However, neither EndFVs nor any solo LTRs were present in other bird species from same order (Pelecaniformes), including the little egret (*Egretta garzetta)*, crested ibis (*Nipponia nippon*) and Yellow-throated sandgrouse (*Pterocles gutturalis)*. Clearly, the study of additional genomes sampled across the avian phylogeny is merited.

### Estimation of ERV insertion times

To approximately estimate the insertion time of endogenous foamy viruses in two birds, we utilized the LTR-divergence based on the degree of sequence divergence between the 5’ and 3’ LTRs with a known host nucleotide substitution rate (Dangel, et al. 1995; Johnson and Coffin 1999). Accordingly, the two intact pairwise LTRs of ERVs-Spuma-Cma were selected for time estimation (Supplementary Table S5). This analysis revealed insertion times of 3.15 and 13.95 MYA (million years ago), close to the estimated time of common ancestor of Maguari Stork and Oriental Stork (Jetz, et al. 2012). However, the presence of multiple premature stop codon suggests the invasion was ancient, and all estimates of integration times should be treated with caution (Kijima and Innan 2010). Clearly, the discovery of additional EndFVs will shed more light on the time-scale of these retroviral integration events. In sum, we describe the presence and evolution of two novel avian endogenous foamy viruses. This discovery fills the last major gap in our understanding of the taxonomic distribution of the foamy viruses and helps reveal the evolutionary interactions between retroviruses and their hosts over extended time-periods.

## Supporting information

Supplementary data set S1

Supplementary data set S2

## Acknowledgements

This work was supported by the Special Key Project of Biosafety Technologies (2017YFC1200800) for the National Major Research & Development Program of China, and Strategic Priority Research Program of the Chinese Academy of Sciences (XDA13010500). J.C. is supported by National Natural Science Foundation of China (31671324, 31970176) and CAS Pioneer Hundred Talents Program. E.C.H. is supported by an ARC Australian Laureate Fellowship (FL170100022). G.Z. is supported by Strategic Priority Research Program of the Chinese Academy of Sciences (XDB31020000) and Villum Foundation. We thank B10K consortium for the early access of all bird genomes produced by B10K.

