## Supplementary Fig.S1-S3; Supplementary Table S1-S5 for "Identification and evolution of avian endogenous foamy viruses"

**Chen et al**

**The file includes**

- **Fig. S1.** Detailed descriptions of the putative genomes of ERV-Spuma.1-Cma and ERV-Spuma.2-Cma and ERV-Spuma.1-Cbo.
- **Fig. S2.** Conserved domain alignments of the original ERV-Spuma.1-Cma proteins and foamy virus.
- **Fig. S3.** Comparison of two orthologous endogenous foamy viruses discovered in two bird species.
- **Table S1.** Information on the 147 bird genomes in Genbank used for data mining.
- **Table S2.** Information on the representative retroviruses.
- **Table S3.** The significant hits identified in *Ciconia maguari* and *Ciconia boyciana* genomes.
- **Table S4.** The endogenous foamy viruses identified in *Ciconia maguari* and *Ciconia boyciana* genomes.
- **Table S5.** Dating the ERVs-Spuma-Cma insertion based on LTR-LTR divergence.
- **References**

### ERV-Spuma.1-Cma

**| 5′LTR start**

1 **TG**TCGCAGGGCACAGAGCAGCCCTTGGTGGAGTAAGGAGCGGTCTGACTGCCGTCAAAGCCTCCCATGAACAAATAACAAATAGGGATAG

91 GCTGCTAAACCACAAAGGCAGACTCACATATGCTTGCTTGCTTAAGTAGGTTTGAATCCCTGAAATAGGGGATTTCCTGTTTGAACGTCA

181 CTCGGTACATAAACATCGAACAAGTTCTGGTATTATGCTGCCTATGTAAAAGATGAAGGTGTGTAAAAGTAAGATCGAGTGTAAAGCCAG

271 CAGATTTCCAGCCTATTGGCACTTGGTTGCGTCCAAGCAAGGAGACCTCTGGCTCGGTATGTAAATCTTATTACTTAATGCTTGCTCAGG

361 TCACTTAGTATTTTATTATTAATTGTGTCTGATAGCTCTGTATGATGTGTATATAGTAATAAATAGTAATTAATATATCTGCTCCTTGTG

451 GCCCTGTAATTTATTGTCAGGGGCTTAATATAAGCAACCTGTGCATAATCCCAACTATTCCTGATTAGCCCAGGCCAAGTAGTGGAATAA

**5′LTR end |**

541 GTGATCTACGTGAGGCCCATACCCGTGA**CA**AT**TGGTGCCCAACGTGGGGC**TCAAGGAGCACGATATATAACTGGTACCCTTAGAATAAAT

**| Gag start** **PBS**

M A H N P Y N L E S F N Q R V Q M L Y E R P P T H G

631 CCTAGATCTAGGTATGGCTCATAATCCCTATAATTTAGAGAGTTTTAATCAGAGGGTTCAGATGTTGTATGAACGCCCTCCAACACATGG

E N I T V C I M N G P W G I G D R Y K R K R L E L Q D Q G G

721 TGAAAATATCACCGTATGCATTATGAATGGTCCGTGGGGAATTGGTGATAGATATAAACGAAAAAGGTTAGAATTGCAAGATCAAGGAGG

A N L P I P Q W Q H Q D G R I E R T E I V I H A N F A E T L

811 GGCAAATTTGCCTATCCCCCAGTGGCAACACCAGGATGGACGAATAGAAAGAACTGAAATTGTAATTCATGCAAATTTTGCTGAAACATT

N W L G Q P P D I N T G E D R H G P M A H E P F T P G D E I

901 AAATTGGTTAGGACAACCACCGGACATTAATACTGGAGAGGATCGACATGGGCCAATGGCTCATGAACCTTTTACTCCAGGAGATGAGAT

C E G Y L P I T L E E L E Q L N Q P A N L R I E A A L L A R

991 TTGTGAAGGATATTTACCTATAACCTTGGAAGAATTAGAGCAACTAAACCAACCAGCAAATTTGAGAATAGAGGCAGCATTATTAGCAAG

L Y S Q Q R T Q R R T N V G T G G A R A P L I P V Q V T S L

1081 ATTATATAGTCAACAAAGGACACAAAGAAGGACAAATGTTGGAACGGGAGGAGCTAGAGCTCCATTAATACCAGTTCAAGTTACATCTCT

P F S N I R A A V G P T P M D I K N I F S W V A E R I N V L

1171 CCCCTTTAGTAATATACGAGCAGCAGTAGGACCCACTCCCATGGATATTAAAAATATATTTTCCTGGGTGGCTGAAAGGATTAATGTATT

E G V L P H M N N A A R R Q V V N S L V P Y Q L S L N E E E

1261 AGAGGGGGTATTGCCCCACATGAATAATGCAGCCAGACGGCAGGTGGTTAACTCACTAGTTCCCTATCAATTATCATTAAATGAAGAGGA

C V S W D Q M I S C L Y S K A H G H I P T A K L G E E L Q R

1351 ATGTGTATCTTGGGATCAAATGATATCATGTTTATATAGTAAAGCTCATGGACATATTCCTACTGCTAAATTAGGAGAGGAATTACAGAG

I S S E Q G I K T A F Q L G L A M T N Q N Y G H V W R I I K

1441 AATAAGCAGTGAACAAGGTATAAAGACAGCGTTCCAATTAGGATTAGCTATGACAAATCAGAATTATGGACATGTGTGGAGAATCATTAA

N L V P G Q A P L A E I T R R L E A L P S D Q E R I R Q F Q

1531 AAATCTAGTTCCCGGTCAGGCTCCCCTAGCAGAAATTACTCGCAGATTAGAAGCATTACCAAGTGACCAGGAACGCATAAGGCAATTTCA

T I V D M V Y R M L D L D P T G R R T G T S R A S P V N P S

1621 AACTATAGTGGATATGGTCTATCGAATGTTAGACCTTGATCCTACAGGAAGAAGGACAGGAACATCAAGGGCTAGTCCAGTGAATCCCTC

P S S Q G K S K G K K P F M K V F P K E A P V I P P T F K P

1711 TCCCTCTTCTCAGGGTAAGAGTAAAGGCAAGAAGCCTTTTATGAAGGTATTTCCAAAAGAGGCTCCAGTTATACCTCCAACATTCAAGCC

F K P E E R R Y P A S Q G R T K P R K G G Y N L R P R V T P

1801 TTTTAAACCAGAGGAGAGAAGGTATCCAGCATCTCAAGGAAGAACAAAACCTAGGAAGGGAGGTTATAACCTTAGACCCAGGGTTACTCC

**Gag end |**

P P Q Y G Q W D Q S L G S N R P G P S A P P P R I R *

1891 TCCTCCACAATACGGCCAATGGGATCAGTCTTTAGGATCCAATCGCCCTGGGCCTTCTGCCCCCCCGCCAAGGATAAGATGACAGGTCCT

**| Pol start**

M K V L V Q L Q G Q I I E A

1981 TCAGGAATAAGGAAGAAAATTAACAAGATTAATAACAATAGAACAGACAATGAAAGTTCTGGTCCAGTTACAAGGACAGATAATTGAGGC

E W D S G S E I M I L P K E V L K G L L P I K I I K L I T I

2071 AGAATGGGACTCAGGATCAGAAATAATGATATTACCTAAAGAAGTGTTAAAAGGTCTCTTGCCTATTAAAATAATTAAACTAATTACCAT

T G E I E V S V F Y T T I V I D G K T E G Y E L A E S P D G

2161 TACTGGGGAAATTGAGGTATCTGTATTTTACACAACAATTGTAATTGATGGGAAAACAGAAGGATACGAACTGGCTGAATCCCCTGATGG

**Aspartic protease**

Q A L I S A K D T P W I G V T R K E I E L T I R I D I E K Y

2251 ACAAGCTTTAATTTCAGCCAAAGATACGCCTTGGATAGGTGTAACTAGGAAGGAAATAGAATTAACTATTAGAATTGACATTGAGAAGTA

R E K F R N K P I C Q I K G K S N G N T F W I V L P L V A E

2341 CAGAGAGAAATTCCGAAACAAACCGATTTGTCAGATCAAGGGAAAAAGCAATGGGAACACATTTTGGATCGTTTTGCCCCTTGTGGCAGA

T E N Q V G H R S V L P H N I A A G K I K P K P Q K Q F K I

2431 AACGGAAAATCAGGTAGGACATAGGTCAGTTCTGCCTCATAACATTGCTGCAGGGAAAATAAAACCGAAACCACAGAAACAGTTTAAAAT

N P Q A I P S I Q I V I N Y L L K Q G I P R Q E T S E M N T

2521 AAATCCTCAAGCTATACCGTCTATACAAATAGTTATAAATTACTTACTCAAACAAGGCATCCCCCGGCAAGAAACATCTGAAATGAATAC

P V Y P V P K G E G K W R L V L D Y R A V N K V T P A I A A

2611 ACCAGTTTATCCAGTACCAAAAGGAGAGGGAAAGTGGAGGTTAGTACTAGATTACAGAGCCGTGAATAAAGTAACCCCAGCCATAGCTGC

Q S C H S T G I L M Q L T R K K Y K T T L D L S R W V F G P

2701 ACAAAGTTGTCATTCAACTGGTATTTTAATGCAATTGACACGGAAAAAGTATAAAACAACACTAGACCTCTCTAGATGGGTTTTTGGGCC

**Reverse transcriptase domain**

I L S P K I A F T W C G K T T C L D S A A T R I S K L T S P

2791 CATCCTATCACCAAAGATAGCCTTTACTTGGTGTGGCAAAACAACATGTCTGGACTCAGCTGCCACAAGGATTTCTAAACTCACCAGCCC

I F C R C G G F I K R N P R R L S I C E C I Y F L H D T E K

2881 TATTTTCTGCAGATGTGGTGGATTTATTAAAAGAAATCCCAGACGTCTCAGTATATGTGAATGTATATACTTCTTACATGATACTGAGAA

E H L K T T R H L H N I E R G Q I H S F L K K S E I G K R E

2971 AGAACATCTGAAAACCACAAGACATTTGCACAATATTGAAAGGGGCCAGATACATAGTTTCCTTAAAAAATCTGAAATAGGTAAGCGTGA

V N F L G F A I T N E G R D L T D Q Y E E K L L N L Q P P K

3061 AGTAAACTTTTTGGGCTTTGCAATTACTAATGAGGGAAGAGATCTAACAGATCAATATGAAGAAAAACTACTCAATTTACAGCCCCCAAA

T L K Q L Q S I L G F L N F A H P F I S N F A E L V K S L H

3151 GACTCTTAAGCAACTGCAGAGTATTTTAGGATTTTTGAATTTTGCCCATCCTTTTATTTCCAATTTTGCAGAATTGGTAAAATCTCTCCA

D A I I K A N N N E S F W E R N Q D A L D D L I T A I K Q A

3241 TGATGCCATCATTAAGGCAAATAACAATGAATCCTTTTGGGAACGAAACCAAGATGCTTTAGATGATTTAATCACAGCAATTAAACAAGC

A L L T E R D P T K P L A V K L H V S P E G Y I I A F S R R

3331 AGCTCTTTTAACTGAAAGAGACCCAACCAAACCCCTAGCAGTAAAGCTACACGTGTCTCCTGAAGGCTATATAATAGCCTTCTCCCGAAG

L Y V R L Y N M A D R F P F Q Y T S I I F K R A E K R F T L

3421 GCTGTACGTCAGGCTATATAATATGGCAGATAGATTTCCCTTTCAATACACTTCTATAATATTTAAGCGAGCCGAAAAAAGATTCACTCT

T E K L L T V M Q Y A L I K S F D I A Q G Q M I H V Y S P L

3511 GACTGAAAAACTCTTGACTGTAATGCAATATGCTCTAATTAAAAGTTTTGACATAGCTCAGGGACAAATGATTCATGTTTACTCTCCATT

R C P E T L Q R H T I P E R K V L S S R W L K W M S H I E N

3601 ACGATGCCCTGAGACATTACAAAGACATACCATCCCTGAAAGAAAAGTTTTATCATCACGATGGCTAAAGTGGATGAGTCATATAGAGAA

P Q I K F H Y D E E L P D L A S L Q T P S E K Q V R I R P L

3691 TCCACAAATCAAATTTCATTACGATGAAGAATTACCTGACTTGGCATCTTTACAAACCCCCAGTGAGAAACAGGTCAGGATCAGGCCATT

T E Y K Q I Y Y L Y G S A T T N E H K R Q A R M D A V Q A I

3781 GACAGAATATAAACAAATATATTACCTCTACGGAAGTGCAACAACTAATGAGCATAAAAGGCAGGCCAGAATGGATGCAGTACAAGCCAT

F N P D Y Q V L N V W S I P L G Q H L A Q Y A E V A A L E F

3871 TTTTAATCCAGACTACCAAGTTCTAAATGTTTGGAGTATTCCATTAGGGCAGCATCTAGCGCAGTATGCAGAAGTGGCGGCACTAGAATT

A L Q Q I P M D Q T P R L I I T D S D Y V S K R Y N S K L E

3961 TGCCCTTCAACAAATCCCGATGGACCAAACCCCAAGATTAATTATTACAGATTCAGACTATGTCTCTAAACGTTATAATTCTAAGCTAGA

**RNaseH domain**

F W E S N G F C N A K G K P L H H I S L W K S I S E L K K I

4051 ATTTTGGGAATCTAATGGGTTTTGTAATGCAAAAGGCAAACCATTGCACCATATCTCATTATGGAAAAGCATTTCTGAACTAAAGAAAAT

K P W V H V T H E P G H R C I G T S V R R A G N A A A D S L

4141 TAAGCCCTGGGTGCACGTCACCCATGAGCCAGGGCATCGATGCATCGGGACCAGTGTACGCAGAGCTGGAAATGCAGCAGCTGATTCACT

A K K A S M I N R V H T E P T I D T D L G Q C I N Y L L Q T

4231 GGCAAAGAAAGCCAGTATGATTAACAGGGTACATACCGAGCCAACAATAGACACCGACCTCGGGCAATGCATTAATTACTTACTCCAAAC

H Q D I K K N Y K C H K D D Q G I Y W I T K P E G E F Q I P

4321 CCACCAGGATATAAAAAAAAATTATAAATGTCATAAAGATGACCAGGGAATATATTGGATTACTAAACCTGAGGGGGAATTTCAAATACC

P T T E R H M I T E R A H P S L G H A H F G R D A T L A V L

4411 CCCCACTACTGAACGACATATGATAACTGAAAGAGCTCATCCCTCTTTGGGACATGCACATTTTGGAAGAGATGCCACCTTAGCAGTTCT

K R K C W W P Y M I Q T V Q Q V L Q Y C S K C V T Y N S A N

4501 GAAACGAAAGTGCTGGTGGCCATACATGATCCAGACAGTACAGCAAGTTTTGCAGTATTGCTCTAAATGTGTCACATATAATTCAGCTAA

R A P I P H D K R T I P E S P F D I L F I D Y I G P L P K C
4591 TCGGGCACCCATCCCCCATGATAAAAGAACTATCCCTGAGTCTCCTTTTGATATATTGTTTATTGATTATATAGGACCATTACCAAAATG

P G Q L D Y V L V I I D G A T S F V W L Y P T T G P T A Q A
4681 TCCTGGCCAATTGGATTATGTTCTGGTGATTATAGATGGGGCAACTAGTTTTGTATGGCTGTACCCCACTACAGGACCAACAGCCCAAGC

**Integrase core domain**

T V R A L T D F C K I A I P K K I H S D Q G P A F T A D I S
4771 CACAGTGCGAGCCCTGACTGATTTTTGCAAGATTGCTATTCCCAAGAAAATACACTCGGACCAAGGTCCTGCCTTTACGGCAGATATAAG

K E F A K K Y N I Q W E Y S T P Y H P Q N S G K V E R A N G
4861 CAAAGAGTTTGCAAAGAAGTACAATATACAATGGGAATACAGCACACCCTACCACCCCCAGAATAGTGGGAAGGTTGAGAGGGCGAACGG

E V K A A L T K L S G S C P G K Q Y A Y I L L V Q L G F N N
4951 GGAAGTCAAAGCAGCTCTCACAAAGTTATCGGGAAGTTGCCCTGGCAAGCAGTATGCCTACATTCTGCTAGTACAATTAGGCTTCAATAA

R P R P S I K R T P F K L L F G V P M N V E F N L T S D L S
5041 TAGACCTAGACCATCTATCAAAAGAACACCATTCAAATTACTCTTTGGGGTACCAATGAATGTAGAATTTAATCTTACTTCAGACCTCTC

R E E Q L P L L A E I Q Y T L A T T S T S P P P C S P H S W
5131 CAGAGAAGAACAATTACCACTCCTAGCAGAAATACAATATACATTGGCGACTACCTCCACATCCCCCCCACCATGCTCACCTCATTCATG

H P L V G L L V Q E R V T T R R P L R P R W K P P T P I I K
5221 GCATCCTCTGGTTGGCCTTCTCGTCCAGGAGAGGGTAACTACTCGACGACCACTACGACCTCGATGGAAACCACCAACTCCGATTATTAA

V L S D R V V E I V D K K G N L K Q G S I D N L K V T P H Q
5311 GGTACTATCTGATCGAGTTGTTGAAATTGTGGACAAGAAGGGCAACCTGAAACAAGGATCAATAGATAATTTAAAGGTAACTCCACATCA

Q Q N G C V T G P G D G V G E G D Q M E K H L T Y V P S D K
5401 GCAGCAGAATGGCTGTGTCACTGGACCAGGGGATGGCGTGGGAGAGGGAGACCAGATGGAAAAGCACTTAACTTACGTACCATCAGACAA

**Pol end |**

T E Q A K I L F Q N Y N E E S D E K I T *
5491 GACAGAACAAGCTAAAATACTGTTCCAAAATTATAATGAAGAATCGGATGAAAAAATAACATAGGCAACATGATGCAGATATTCGGGCTA

**| Env start**

M W I I F T L L I L M T M L G V T C T V
5581 TGCTGCGTATGCTACTAGTACTAGGATAGCAATGTGGATTATATTTACCTTACTCATATTAATGACTATGTTAGGAGTAACTTGTACAGT

I V R L Q W K Y A I E R Q G P T I T W N Q T I H Q S P S I H
5671 AATAGTTAGGTTACAATGGAAATATGCTATAGAGAGACAAGGCCCTACAATTACTTGGAATCAAACAATACATCAATCACCATCTATTCA

R I R R G L Y D H P L L V N V P I T G L K Q G L Y R E P F P
5761 TAGAATCAGGAGAGGATTATATGATCATCCATTACTGGTAAATGTGCCTATTACAGGATTAAAACAGGGACTATACCGAGAACCATTTCC

K P I V A K E R V L G I S Q I L I L D S D H M A E A N N L G
5851 GAAACCGATAGTGGCCAAGGAGAGGGTGCTAGGCATATCCCAGATATTGATATTAGATTCGGATCATATGGCTGAAGCAAATAATTTAGG

H P V G K E I L T Q L L N E I P F E I P L D G P Q T Q Q E Y
5941 ACATCCTGTTGGAAAAGAAATTCTTACTCAGCTCCTAAATGAAATACCTTTTGAAATACCTTTAGATGGTCCACAAACTCAGCAAGAGTA

M Q K K R C H E F A H Y Y W I D Y K E Q R K W P E S Q V I A

6031 TATGCAAAAGAAACGTTGCCATGAATTTGCACATTATTACTGGATTGATTATAAGGAACAACGAAAATGGCCAGAATCCCAAGTAATAGC

D H C P H H G R G Y P R F A S E D Y W V E S P F S V T K Q E

6121 TGATCACTGTCCACACCATGGAAGAGGATACCCTCGATTTGCTAGTGAAGACTACTGGGTAGAATCTCCATTCTCTGTTACGAAACAGGA

W K G Y K G Q N R E Q G Y R H I G Y Q G D V R H L L E H S C

6211 ATGGAAAGGCTACAAAGGTCAAAACAGGGAGCAAGGCTACAGACATATAGGTTACCAGGGGGACGTGAGACATTTACTGGAGCACTCGTG

M D D A Y H S L W N K N K M T Q K E L F L A Y V T Q I K K L

6301 TATGGATGATGCCTACCATTCGTTGTGGAACAAAAATAAAATGACACAGAAAGAGCTATTCTTGGCCTATGTCACTCAGATAAAGAAGCT

I Q D M K T G K L K K D A L L N D W H D Q G K G R W F K P M

6391 GATTCAAGATATGAAAACAGGGAAGTTAAAAAAGGATGCTCTTCTGAATGATTGGCATGATCAAGGAAAAGGGAGATGGTTCAAACCCAT

D D L S F C R H P E L T V F L N G T Y H K H S C M E G D C Q

6481 GGATGACTTGTCATTTTGTAGACATCCGGAACTTACAGTATTTCTGAATGGCACATATCATAAACACTCATGCATGGAGGGTGACTGTCA

M T R A N I T H I K G C K K P N K F T Q T P I C M S I L Q I

6571 GATGACTCGGGCAAATATAACGCACATAAAAGGTTGTAAAAAACCTAACAAGTTCACACAAACACCCATATGCATGTCAATTTTACAGAT

D S K C N W R R F S P T V I L C Q R Y L L Y P K Y S R M E E

6661 TGATTCAAAATGCAACTGGAGAAGATTTTCTCCAACTGTAATATTATGTCAGCGATATTTATTATACCCTAAATATAGCAGGATGGAAGA

V S R G V D T G L L L H D H T F P G F W C I E A K Q V Q R Q

6751 AGTCAGTCGTGGAGTAGATACGGGACTCCTTTTACATGATCACACATTTCCCGGATTCTGGTGCATTGAAGCTAAGCAGGTCCAGCGACA

N Y S L Y S L Y Q Q C L F N S Q K H P V D D V I S G M K Q R

6841 GAATTATTCACTGTATTCTCTATACCAGCAATGTTTGTTTAACTCACAAAAACATCCAGTGGATGACGTAATTAGCGGAATGAAGCAGAG

L S V Q K Q E E G Y P C N L T T C Q P V S I L D I S G G Q A

6931 ATTATCTGTACAAAAACAGGAAGAAGGGTATCCGTGCAACTTAACCACATGCCAGCCAGTCTCAATCTTAGACATATCAGGAGGACAAGC

I W G T N E T F L N Y T I V D T P K K P R G S Y T K R K R S

7021 TATCTGGGGCACAAATGAAACCTTCCTAAATTACACGATAGTAGACACACCTAAAAAACCAAGAGGCAGCTATACTAAGCGAAAGAGATC

T T N L Q K I Q E A G L I P D S S I T K T A K I S D L N D K

7111 TACTACTAATCTACAAAAGATACAAGAGGCCGGACTGATCCCAGATAGTTCCATTACTAAAACTGCCAAAATATCTGATTTAAATGATAA

Q L A K G L H L L R D H L I T F P E H T I D V I Q M S Q S I

7201 ACAATTAGCAAAAGGGTTACATCTGCTACGAGATCACCTTATTACATTTCCTGAACATACAATTGATGTGATTCAAATGAGTCAATCCAT

I A V M T H S H I Q N L R I L L T E G K V D W D T L N S T W

7291 TATCGCAGTAATGACACATTCTCATATACAGAATTTAAGAATCCTTTTAACTGAAGGCAAGGTTGATTGGGACACTTTAAACTCAACCTG

I Q E Q L R V S D E M M T L I R R T A R G L A Y D I Q Q R V

7381 GATACAAGAACAATTACGAGTCTCAGATGAAATGATGACTTTAATAAGAAGAACAGCCAGGGGATTGGCTTATGATATACAACAGCGGGT

D K P E K G V W E I S L Y Y E I V I P R R I Y S T N W K I I

7471 TGACAAACCAGAAAAAGGAGTCTGGGAAATCTCCCTATATTATGAAATTGTTATACCACGAAGAATATATTCTACTAATTGGAAAATTAT

N Y G H L V Y T G N R G G R V W L K H P C T L I T Q G C G E

7561 CAATTATGGTCATCTGGTATATACTGGAAATAGAGGAGGAAGAGTCTGGTTAAAACATCCCTGTACCCTAATAACCCAAGGGTGTGGTGA

V K Y L E V R E C Y E Q D Y L I C D E V I K H E P C G N Q T

7651 AGTAAAATATTTAGAAGTAAGAGAATGCTATGAACAAGATTATCTTATTTGTGATGAGGTCATCAAGCATGAGCCATGTGGTAACCAAAC

G S R C P I M V E P I V S P Y L R I G P L K N G N Y I V T T

7741 TGGATCCAGATGTCCTATAATGGTAGAACCAATTGTTTCCCCATACCTTAGAATTGGCCCATTGAAGAATGGGAATTACATAGTGACGAC

S L D E C S I P P Y Q P S L I T V N E T V T C Y G Y E F K S

7831 CAGTTTGGATGAGTGTAGCATACCACCATACCAGCCATCTCTGATCACAGTAAATGAGACAGTGACTTGCTATGGATATGAGTTTAAATC

P L R I N T V I Q E N I H M P P L S V S L P H M I G I I A D

7921 ACCCCTTAGAATAAATACCGTAATACAAGAAAATATACATATGCCACCACTGTCAGTCAGCTTACCGCATATGATAGGTATTATAGCTGA

L R Q I K I E L A S S W D S V H D V T E R A N T E L L R I D

8011 TCTTAGACAAATAAAAATCGAGCTGGCTTCCTCTTGGGACTCAGTTCATGATGTGACTGAGAGAGCAAATACTGAACTACTGAGAATAGA

L R E G Y T P Q W L N R L S E S I A D I W P A A A G A I K G

8101 TTTACGTGAAGGATATACACCTCAATGGTTAAATAGACTGTCGGAATCCATTGCAGACATCTGGCCTGCTGCAGCTGGAGCTATCAAGGG

**Env end |**

I A N G I K D L T S G N L V Q L L I C *

8191 CATAGCTAATGGAATCAAGGATCTAACATCTGGAAATTTGGTACAGCTTTTGATTTGTTAGCTTATGCAAACCCCGTGATAATAGGAGTG

8281 GTTTTGTTGATTACATTAGTGCTAGTCATCAAATTAATCAGTTGGATTGCTACGAAACACAAAACAGTGAGAAGATGGAATCACAAGATC

8371 AGGCTATGTTAGCAATGTTATGCCTTTATAGACCTCTTACTGTAAAAAGTATACTTATTATAAAACTATCCTGGTACCAATGCCGAGTTG

**Internal Promoter**

8461 TGACCAGGAGTCTACTTTTATTTTTAGATATACTTGTCATTATTATGCTTATATCTATCGTAGTAAATATCTAATTAAGCCAAGAGAAAA

**| ORF1 start**

M E T E H T T P L L Q S D N S N I G D G I L S V H T L K P D

8551 TGGAGACAGAGCATACTACACCACTACTCCAAAGTGATAACTCAAACATAGGGGATGGAATCCTATCTGTGCACACATTAAAACCGGACA

T K I D H G Q Y P V V M Q Q T L T I L K G A P P V W I H T L

8641 CCAAAATTGATCATGGACAGTACCCTGTGGTCATGCAACAGACCCTTACTATTCTCAAAGGTGCACCCCCTGTCTGGATACACACGCTGG

A K E L A L I T L S D G T E I L P N H M G W P I P M E Q P Q

8731 CGAAGGAACTAGCACTGATCACACTATCCGATGGGACGGAGATACTGCCCAACCACATGGGATGGCCAATCCCGATGGAACAGCCACAAC

H L L Y F Q S L H H W M Y C S S L R N W V Q T L I T T H C L

8821 ATCTCCTATACTTTCAGAGTCTTCATCATTGGATGTACTGCAGCTCTTTGAGGAACTGGGTGCAGACCTTGATCACCACTCATTGTCTGA

Nonsense mutation (C→T)    Large insertion (CGATAACATATTCAAAGA)

N L I F P R L G Q V Y W F L T L T C C N I A S G L T H G R C

8911 ATCTGATATTCCCTCGCCTGGGACAGGTTTATTGGTTTTTAACGTTGACATGCTGTAACATAGCTAGTGGATTGACACATGGACGTTGTT

Insertion (A)

Y M G L W E P G N N G T A E G W L F L G W K Q C L Q A Q G T

9001 ACATGGGTCTTTGGGAACCTGGAAACAATGGGACTGCTGAGGGATGGCTGTTCCTGGGTTGGAAACAATGCCTACAAGCACAAGGTACAA

S N L W C H K P L L L P M R C Y D T N F A F V S N E S I Q L

9091 GTAATCTTTGGTGCCATAAGCCATTATTACTACCCATGCGTTGTTATGATACTAACTTTGCATTTGTTAGCAATGAGAGTATCCAGTTGA

**ORF1 end |**

K T V T C V C Y G I K A I T Q I L V R P L G I L D I N I *

9181 AGACTGTAACGTGTGTGTGCTATGGTATAAAAGCTATAACTCAAATTTTAGTAAGACCCCTTGGGATATTAGATATCAACATCTAGCAAT

9271 GTTAAAAAAAATGGATGTACTAATGATATACAATAATCTTGTACTAACTGTCAAAGAAAATTGCTATACGAATGATTACTAAGAACCAAA

**| 3′LTR start**

9361 AAGATAATTGGCAGGCTGTG**ACGGCAAGGAGAGGGT****G**TCGCAGGGCACAGAGCAGCCCTTGGTGGAGTAAGGAGCGGTCTGACTGCCGTC

**PPT**

9451 AAAGCCTCCCATGAACAAATAACAAATAGGGATAGGCTGCTAAACCACAAAGGCAGACTCACATATGCTTGCTTGCTTAAGTAGGTTTGA

9541 ATCCCTGAAATAGGGGATTTCCTGTTTGAACGTCACTCGGTACATAAACATCGAACAAGTTCTGGTATTATGCTGCCTATGTAAAAGATG

9631 AAGGTGTGTAAAAGTAAGATTGAGTGTAAAGCCAGCAGATCTCCAGCCCATTGGCACTTGGTTGCATCCAAGCAAGGAGACCTCTGGCTC

9721 GGTATGTAAATCTTATTAATGCTTGCTCAGGTCACTTAGTATTTTATTATTAATTGTGTCTGATAGGTCTGTATGATGTGTATATAGTAA

9811 TAAATAGTAATTAATATATCTGCTCCTTGTGGCCCTGTAATTTATTGTCAGGGGCTTAATATAAGCAACCTGTGCGTAATCCCAACTATT

9901 CCTGATTAGCCCAGGCCAAGTAGTGGAATAAGTGATCTACGTGAGGCCCATACCTGTGA**CA**

**3′LTR end |**

### ERV-Spuma.2-Cma

| 5′LTR start

1 **TG**TTGCAGGGGACAGACCAGCCCTTGGTGGAGTAAGGAGCAATCTGACCACCGTCAAAGCCTCCCATGAACAAATAACAAATAGAGATAG

91 GTTGCTAAACCGCAAAGGCAGACTCACATATGCTTGCTTCCTAAAGGAAGTTTGAATCCTTGAAACAGAAGATTTCCTGTTTGAACACCA

181 CTCGGTACATAAACATCAAAAAAGTTCTGGTATTACGCTACCTACAGAAAAGATGAAGGTGTATAAAAGTAAGATCGAGTGTAAAGCCAG

271 CAGATCTCCAGCCCATTGGCACTTGGTTGCATCCAAGCAAGGAGACCTCTGGCTCAGTANATTGGCACTTGGTTGCATCCAAGCAAGGAG

361 ACCTCTGGCTCAGTACGTAAATCTTATTAGTTAATGCTTGCTCAGGTCACTTAATATTTTATTATTCATTGTGTTTGATACTTCTGTATG

451 ATGTGTATATGGTAATAAATAGTAATTAATATATCTGCTCCTTGTGGCCCTGTAATTTATTGTCAGGGGCTTAATGTAAGCAACTTGTGC

5′LTR end |

541 GTAATCCCAACTATTCCTGATTGGCCCAGGCCAAGTAATGGAATAAGTGATCTATGTGAGGCCTATACCCACAA**CA**AT<u>TGGCGCCCAATG</u>

PBS

631 <u>TGGGC</u>TCTAGGAGCACAATATATAACTGGTACCCTTAGAATAAATCCTAGATCTAGGCATGGCTCATAATCTGTATAATTTAGAGAGTTT

| Gag start

M L Y Q R P P R H G E N I T I R I T N G P W R T G

721 TAATCAGAGGTTCAGATGTTGTATCAACGCCCTCCAAGACACGGTGAAAATATCACCATACGCATTACGAATGGTCCATGGAGAACTGGT

D R Y T Q I T L E L Q D Q G G T N L P I P E L A T L D G Q I

811 GATAGATATACACAATAACTTTAGAACTGCAAGATCAAGGAGGGACAAATTTGCCTATCCCCGAGCTAGCAACACTGGACGGACAAATA

E R T E I V I H A N F A E T L N W L G Q P P D I N T G V D R

901 GAAAGAACTGAAATTGTAATTCATGCAAATTTTGCTGAAACGTTAAATTGGTTAGGCCAACCACCAGACATTAATACTGGAGTGGATCGA

H G P M A H E P F P P R D E I C E G Y L P I T L E E L E Q I

991 CATGGACCAATGGCTCATGAACCTTTTCCTCCAAGAGATGAGATTTGTGAAGGATATTTACCTATAACCTTGGAAGAATTAGAGCAAATA

N Q P G N L R I E A A I L A R L Y S Q Q R T Q I G T N V G T

1081 AACCAACCAGGAAATTTGAGAATAGAAGCAGCAATATTAGCAAGATTATATAGCCAACAAAGAACACAAATAGGGACAAATGTTGGAACG

G G A R I P L V L V Q V A S L P F G N I R A A V G H G Y K K

1171 GGAGGAGCTAGAATTCCATTAGTACTGGTCCAAGTTGCATCTCTCCCCTTTGGTAATATACGAGCAGCAGTGGGACATGGATATAAAAAG

I F S W M A E R I N I L E G V L P N M N N A T R C Q L V N S

1261 ATATTTTCCTGGATGGCTGAGAGGATTAATATATTAGAGGGGGTATTGCCCAATATGAATAATGCAACCAGATGTCAGCTGGTTAACTCA

L V P Y Q L S L N E E E C L S W D Q I I S C L Y A K A H R H

1351 CTAGTTCCCTATCAATTATCATTAAATGAAGAGGAATGTTTATCCTGGGACCAAATTATATCATGTTTATACGCTAAAGCTCATAGACAT

I P T A K L G E E L Q R I S S E Q G Y K T A F S S G L A M T

1441 ATTCCTACTGCTAAATTAGGAGAGGAATTACAGAGAATAAGCAGTGAACAAGGATACAAGACAGCATTCTCATCAGGATTAGCTATGACA

N Q N Y G H L W G T V K N L V P G Q A P L A E I T C R L E V

1531 AATCAGAATTATGGACATCTGTGGGGAACTGTAAAAAATCTAGTTCCTGGTCAGGCTCCACTAGCAGAAATTACTTGCAGATTAGAAGTG

L P S D R E H I R Q F S T I V D T V Y R M L D L D P T G K M
1621 TTACCAAGTGACCGGGAACACATAAGGCAATTTTCAACTATAGTGGATACAGTCTACCGAATGTTAGACCTTGATCCTACAGGGAAAATG

T G R S R A S P V N P S P S P Q G K S R N K K P F K K R F P
1711 ACAGGAAGGTCAAGGGCTAGTCCAGTGAATCCCTCTCCCTCTCCTCAGGGCAAGAGTAGAAACAAGAAGCCTTTTAAGAAGAGATTTCCA

K E I P V I P P T F K P F K S E E R R Y P A S Q G K T E S T
1801 AAAGAAATTCCAGTTATACCTCCAACATTCAAGCCTTTTAAATCAGAAGAAAGAAGGTATCCGGCATCTCAAGGAAAAACAGAATCTACT

K G A Y N L K P R V T P S R Y G Q W A Q S S A S N C P G P S
1891 AAAGGAGCTTATAACCTTAAACCCAGGGTTACTCCTTCACGATATGGACAATGGGCTCAGTCTTCAGCATCCAATTGCCCTGGGCCTTCT

**Gag end |**

A P L P A P P G I T S Q V L R E *
1981 GCCCCCCTCCCCGCCCCCCCGGGGATAACAAGCCAGGTCCTTCGGGAATAAGGAAGAAAATTAACAGGATTAATAAAAATAGAAAGGACA

**| Pol start**

M K V L L Q I E G E T I E A E W D S G S E I T I L P K E V L
2071 ATGAAAGTTCTGCTCCAGATAGAAGGAGAGACAATTGAGGCAGAATGGGACTCAGGATCAGAAATAACAATATTACCTAAAGAAGTGTTA

K G L L P I K R I K L M T I T G E T E V P V F Y T T I I I D
2161 AAAGGTCTCTTACCTATTAAAAGAATTAAACTAATGACCATTACTGGGGAGACTGAGGTACCTGTATTCTACACAACAATTATAATTGAT

G K K R R I E V A E S P D G Q A L I S V K D T P W I G I T R
2251 GGGAAAAAGAGAAGGATAGAAGTGGCTGAATCACCTGATGGACAAGCTTTAATTTCAGTGAAAGATACGCCTTGGATAGGTATAACCAGG

**Aspartic protease**

L G S R I N Y Q N R H R E D T K R N S E T N Q F V R S R G K
2341 CTAGGAAGTAGAATTAACTATCAGAATCGACATCGAGAAGATACAAAGAGAAATTCTGAAACAAACCAATTTGTCAGATCGAGGGGAAAA

A I G T I F R S F C P L V A E M G K S G R T Q V N S A S T G
2431 GCAATTGGAACAATTTTTCGATCATTTTGCCCCCTTGTGGCAGAAATGGGAAAATCAGGTAGGACACAGGTCAATTCCGCCTCTACAGGG

K I K P K A Q K Q F E
Q K Q F E I N P Q A I P S I Q I V L N D L L K Q
2521 AAAATAAAACCGAAAGCACAGAAACAGTTTGAAATAAATCCTCAAGCTATACCGTCTATACAAATAGTTCTAAATGACTTACTCAAACAA

S V L R Q E T S E M N T P V Y P V P K G E G K R R L V L D Y
2611 AGTGTCCTCAGGCAAGAAACATCTGAAATGAATACACCAGTTTATCCAGTACCAAAAGGAGAGGGAAAGAGGAGATTAGTACTAGATTAT

R A M N K V T P A I A A Q N C H S T S I L M Q L T Q K K Y K
2701 AGAGCCATGAATAAAGTAACCCCAGCCATAGCTGCACAAAATTGTCATTCAACCAGTATTTTAATGCAATTGACACAGAAAAAGTATAAA

**Reverse transcriptase domain**

T T L D L S N G F W A H P I T K D S Q W I T A F T C C G I Q
2791 ACAACACTAGACCTCTCTAATGGTTTTTGGGCCCATCCTATCACCAAAGATAGCCAATGGATAACAGCCTTTACTTGCTGTGGCATACAA

H V W T R L P Q G F L N S Q A L F S A D V V D L L K E F P E
2881 CATGTCTGGACTCGGCTGCCACAAGGATTTCTAAACTCACAAGCCCTATTTTCTGCAGATGTGGTGGATTTATTAAAAGAATTCCCAGAG

V S V Y V D D T Y F S R D T E K E H L K T L Q G I C T I L K
2971 GTCTCAGTATATGTGGACGATACATACTTCTCACGTGATACTGAGAAAGAACATCTGAAAACTCTGCAAGGCATTTGCACAATATTGAAA

G A G Y I V S F K K S E I G N H E V N F L G F V I T N E G R

3061 GGAGCCGGATATATAGTTTCATTTAAAAAATCTGAAATAGGTAACCATGAAGTAAACTTTTTGGGCTTTGTGATTACTAATGAGGGAAGA

G L T D E Y K E K L L N L Q P P K T L K Q L Q N I L G F L N

3151 GGTCTAACAGATGAATATAAAGAAAAACTACTCAATTTACAGCCCCCAAAGACTCTTAAGCAACTGCAGAATATTTTGGGATTTTTGAAT

F A C A F I S N F V E L V K P L H D A I I K A N N N E P F W

3241 TTTGCCTGTGCTTTTATTTCCAATTTTGTAGAATTGGTAAAACCTCTCCATGATGCCATCATTAAGGCAAATAACAATGAACCCTTTTGG

E Q K Q Q D A L D G L I M A I N Q A A L L T E R D T T K P Q

3331 GAACAAAAACAACAAGATGCTTTAGATGGATTAATCATGGCAATTAACCAAGCAGCTCTTTTAACTGAAAGAGACACAACCAAACCCCAG

Q Q K T P F K L L F G V S K K V E F N L T S D L S R E E Q L

3421 CAGCAAAAAACACCATTCAAATTACTCTTTGGGGTATCAAAGAAAGTAGAATTTAATCTTACTTCAGACCTCTCCAGAGAAGAACAATTA

A L L A E I R P M L A T T S T S S P L P S P R S W R P L V G

3511 GCACTCCTAGCAGAAATACGACCTATGTTGGCGACTACCTCCACATCCTCGCCACTACCCTCACCTCGTTCATGGCGTCCTCTGGTTGGC

**Pol end |**

L L V Q E R V A T R R P L *

3601 CTTCTTGTCCAGGAGAGGGTAGCTACTCGACGACCATTATAACCTCAGTGAAAACCACCAACTCCAATTATTAAGGTACTATCTGATCGA

3691 GTTGTTGAAATTGTGGATGAGAAGAGTAACCTGAGACAAGTATCAACAGATAATTTAAAAGTAACTCCACATCAGCAACAGAATGACCTT

3781 GTCATTGCACCAGTGGATGGCATGGGAGAGGGAAACTAGATGTAGAAAAGCACTTAATACCATCAGACAAGACAGAACATCCTACAAATA

**| Env start**

M F Q N Y I E E L D E K I T W K T R C R Y W G Y A A C A T S

3871 ATGTTCCAGAATTATATTGAAGAATTGGATGAAAAAATAACATGGAAAACACGATGCAGATATTGGGGCTATGCTGCATGTGCTACTAGT

A R I A M Q I I F T L L I L M T M L G V T C T V I V R L Q W

3961 GCTAGGATAGCAATGCAGATTATATTTACCTTACTCATATTAATGACTATGTTAGGAGTAACTTGTACAGTAATAGTTAGGTTACAATGG

K Y A I E R Q G P T I T W N Q T I Y Q L P S I H R I R R G L

4051 AAATATGCTATAGAGAGACAAGGCCCTACAATTACTTGGAATCAAACAATATATCAATTACCATCTATTCATAGAATTAGGAGAGGATTA

Y H H P L P V N V T I T G L K Q G L Y W E P F P K L I V A K

4141 TATCATCATCCATTACCAGTAAATGTGACTATTACAGGATTAAAACAGGGACTATACTGGGAACCTTTTCCAAAACTGATAGTGGCCAAG

E R V L G I S Q I L I L D S D H M A E A K N L G D P V G K E

4231 GAGAGGGTGCTAGGCATATCCCAGATATTGATATTAGATTCGGATCATATGGCTGAAGCAAAAAATTTAGGAGATCCTGTTGGAAAAGAG

I L I Q L L N E E M K Q L K D I T L S F E I P L D G P Q T Q

4321 ATTCTTATTCAGCTCCTAAATGAGGAAATGAAACAGCTTAAAGATATCACTCTTAGTTTTGAAATACCTTTAGATGGTCCACAAACTCAG

Q E Y I Q K K C Y H E F A H C Y W I D Y K E Q G K W P E P Q

4411 CAAGAGTATATTCAAAAGAAATGTTACCATGAATTTGCACATTGTTACTGGATTGATTATAAGGAGCAAGGAAAATGGCCAGAACCCCAA

V I A D H C P H P R S G Y P R F S G K N Y W V E S P S S V T

4501 GTAATAGCTGATCATTGCCCACACCCTAGAAGCGGATACCCTCGATTTTCTGGTAAAAATTACTGGGTAGAATCTCCATCCTCTGTTACA

K Q E W E R L Q R L K Q G A R L Q T Y R L P G G H E T F T G
4591 AAACAGGAATGGGAAAGGCTACAAAGGTTAAAACAGGGAGCAAGGCTACAGACATATAGGTTACCAGGGGGACATGAGACATTTACTGGA

A L M C T D D A Y H L W W R K N K T T Q K D L F L A S V T Q
4681 GCACTAATGTGTACAGATGATGCCTACCATTTATGGTGGCGCAAAAATAAAACAACACAAAAAGACCTATTCTTAGCCTCTGTCACTCAA

I K K L I Q D T K T G K L K K V F L L N G W H D Q G K G K W
4771 ATAAAGAAGCTGATTCAAGATACAAAAACAGGGAAATTAAAAAAGGTTTTCCTTCTGAATGGTTGGCATGATCAAGGAAAAGGGAAATGG

F K S M D D L S F C R H P E L T V F L N G T Y Y K H S C T D
4861 TTCAAATCCATGGATGACTTGTCATTTTGTAGACATCCGGAACTTACAGTATTTCTGAATGGCACATATTATAAACATTCATGCACGGAC

G D C Q S T Q A N I T Q I K G C K N L T S S P K H P Y A C Q
4951 GGTGACTGTCAGAGCACTCAGGCAAATATAACACAGATAAAAGGTTGTAAAAACCTAACAAGTTCTCCTAAACACCCATATGCATGTCAA

F Y R L I Q N A T G E D F L Q L Q Y Y D Q R Y L L Y P K Y S
5041 TTTTACAGATTGATTCAAAATGCAACTGGAGAAGATTTTCTCCAACTGCAATATTATGATCAGCGATATTTATTATACCCTAAATATAGC

R M E K V S R G V D T G L L L H D H T F P G P W C V E A K Q
5131 AGGATGGAAAAAGTCAGTCGTGGAGTAGATACGGGACTCCTTTTACATGATCACACATTTCCCGGACCCTGGTGTGTTGAAGCTAAGCAG

V W Q Q N Y S L Y S L Y Q Q C L F H S Q K H P V D D V I S G
5221 GTCTGGCAACAGAATTATTCACTATATTCTTTATACCAGCAATGTTTATTTCACTCACAAAAACATCCAGTGGATGATGTAATTAGCGGA

M K Q R L F V Q R Q G E G Q P Y S L T T C H P V S I L D K S
5311 ATGAAGCAAAGATTATTTGTACAAAGACAAGGAGAAGGGCAGCCGTACAGCTTAACCACGTGCCACCCAGTCTCAATCTTAGACAAATCA

R G R A I W G A N E T F L N Y T I Q T K K S K G C H T K Q K
5401 CGAGGACGAGCTATCTGGGGCGCAAATGAAACCTTCCTAAATTATACAATACAGACTAAAAAATCAAAAGGCTGCCATACTAAGCAAAAG

R S I T N L Q K I Q E A R L I L G S S I T K I A K I S D L N
5491 AGATCTATTACTAATCTACAAAAGATACAAGAGGCCAGATTGATCCTAGGTAGTTCCATTACTAAAATTGCCAAAATATCTGATTTAAAT

G K E L A K G I H L L R D N L I T F A E H T I D D V I Q M S
5581 GGTAAAGAATTAGCAAAAGGGATACATCTGCTACGAGATAACCTTATTACATTTGCTGAACATACAATTGATGATGTGATTCAAATGAGT

P T I T T V M I H S H I Q N L R I L L T E G K V D W N I L N
5671 CCAACCATTACCACAGTAATGATACATTCTCATATACAGAATTTAAGAATCCTTTTAACTGAAGGCAAGGTTGATTGGAACATTTTAAAC

S T W I Q E Q L Q V T D E I M T L I R R T A R G L A Y D I Q
5761 TCAACCTGGATACAAGAACAATTACAAGTCACAGATGAAATTATGACTTTAATAAGAAGAACAGCTAGGGGATTAGCTTATGATATACAA

Q W V N K P Q K G V W E I S L Q Y D I V I P Q R I Y P T N W
5851 CAATGGGTTAACAAACCACAAAAAGGAGTCTGGGAAATCTCCCTACAGTATGACATTGTTATACCACAAAGAATATATCCTACTAATTGG

K I I N Y G H L V Y T G N K G G R I W L K H P Y T L I T Q G
5941 AAAATTATCAATTATGGTCATCTGGTATATACTGGAAATAAAGGAGGAAGAATCTGGTTAAAACATCCCTATACCCTAATAACCCAAGGG

```
     W  G  E  V  R  Y  L  E  V  R  Q  C  Y  E  Q  D  Y  L  I  C  D  E  V  I  E  H  E  P  Y  G
6031 TGGGGGGAAGTAAGATATTTAGAAGTAAGACAATGCTATGAACAAGATTATCTTATTTGTGATGAGGTCATTGAACATGAGCCATATGGT

     N  Q  T  G  S  R  C  P  I  M  V  E  P  I  V  S  P  Y  L  R  T  D  L  L  K  N  G  S  Y  I
6121 AACCAAACTGGATCCAGATGTCCTATAATGGTAGAACCAATTGTCTCCCCATACCTTAGAACTGACCTATTGAAGAATGGGAGTTACATA

     V  M  T  S  L  D  E  C  N  V  P  P  Y  Q  P  S  L  I  A  V  H  E  T  V  T  C  Y  G  Y  E
6211 GTGATGACCAGTTTGGATGAGTGTAACGTACCACCGTACCAACCATCTCTGATCGCAGTACATGAGACAGTGACTTGCTATGGATATGAA

     F  K  P  P  L  R  I  N  T  V  I  Q  E  N  I  P  V  P  P  L  S  V  S  L  L  H  M  I  G  I
6301 TTTAAACCACCCCTTAGAATAAATACAGTAATACAAGAAAATATACCTGTGCCACCACTGTCAGTCAGCTTACTGCATATGATAGGTATT

     I  A  D  L  R  R  I  K  M  E  Q  A  S  S  L  G  L  S  S  C  M  I  K  R  E  N  T  Q  L  L
6391 ATAGCTGATCTCAGAAGAATAAAAATGGAGCAAGCTTCCTCTTTGGGACTCAGTTCATGTATGATTAAGAGAGAAAATACTCAACTACTG

     R  I  D  L  H  E  G  D  I  P  Q  W  L  N  R  L  S  E  S  I  A  D  I  W  P  A  A  A  G  A
6481 AGAATAGATTTACATGAAGGAGATATACCTCAATGGTTAAATAGACTGTCAGAATCCATTGCAGACATCTGGCCTGCTGCAGCTGGAGCT

     I  K  G  I  A  N  G  I  K  D  L  T  S  G  I  F  G  M  A  S  D  L  L  A  Y  A  N  P  V  I
6571 ATCAAGGGCATAGCTAATGGAATCAAGGATCTAACATCTGGAATTTTTGGTATGGCTTCTGATTTGTTAGCTTATGCAAACCCCGTGATA
                                                                                    Env end |
     I  G  V  V  L  L  I  T  L  V  L  V  I  K  L  I  S  W  I  A  T  K  R  K  T  E  *
6661 ATAGGAGTGGTTTTGTTGATTACATTAGTGCTAGTCATCAAATTAATCAGTTGGATTGCTACAAAACGTAAAACAGAGTGAGATGATGGA

6751 ATCACAAGATCAGGCTACGTTAGCAATGTTATGGTTTTACAGAGCTCTTACTGTAAAAAGTATACTTACTCATTATAAAACTATCCTGGT
                                                                             Internal Promoter
6841 ACCAATGCCGAGTTGTGACCAGGAGTCTACTTTTGTTTTTAGACACGTTTGTCATTATTGTGCTGATATCTATCATAGTAAATATCTAAT
                  | ORF1 start
                  M  E  K  E  H  T  I  P  L  P  Q  S  H  N  S  N  I  R  D  G  I  L  S  V  H
6931 TAAGAGAAGAGAAAATGGAGAAAGAGCATACTATACCACTACCCCAAAGCCATAACTCAAACATACGGGATGGAATCCTATCTGTACACA

     T  L  K  P  D  T  E  I  N  H  G  Q  F  P  V  I  M  Q  L  T  L  T  T  L  E  G  A  P  P  V
7021 CATTAAAACCAGACACTGAAATCAATCATGGACAGTTCCCTGTGATCATGCAACTGACCCTTACTACTCTCGAAGGTGCACCACCTGTCT

     W  I  Q  T  P  A  K  E  L  A  L  I  T  L  S  D  G  T  K  I  L  P  S  H  M  G  W  P  V  P
7111 GGATACAGACTCCGGCAAAGGAACTAGCACTGATCACACTATCTGACGGGACTAAGATATTGCCCAGCCACATGGGATGGCCAGTCCCAA

     M  E  Q  P  Q  H  L  L  Y  F  Q  S  L  H  H  W  M  Y  C  S  S  L  R  N  W  V  Q  T  L  I
7201 TGGAACAACCACAACATCTCCTATACTTTCAGAGTCTTCATCATTGGATGTACTGCAGCTCTTTGAGGAACTGGGTGCAGACCTTGATCA

     T  T  H  C  L  N  L  V  F  P  P  L  G  Q  V  Y  W  F  L  T  L  T  C  Y  N  I  A  S  G  L
7291 CCACTCATTGTCTGAATCTGGTATTCCCTCCCCTGGGGCAGGTTTATTGGTTTTTAACGTTGACATGCTATAACATAGCTAGTGGATTGA

     T  H  G  N  C  Y  M  G  L  W  E  S  G  N  N  A  T  A  E  G  W  L  F  W  G  R  K  P  C  L
7381 CACATGGAAATTGCTACATGGGTCTTTGGGAATCTGGAAACAATGCGACTGCTGAGGGATGGCTGTTCTGGGGTCGGAAACCATGCCTAC

     Q  A  Q  G  T  S  N  L  W  C  H  K  P  L  L  L  P  M  H  C  Y  D  T  K  F  A  F  V  S  Y
7471 AAGCACAAGGTACAAGTAATCTTTGGTGCCATAAGCCATTATTACTACCCATGCATTGTTATGATACTAAATTTGCATTTGTTAGCTATG
```

G A C L V E D C N V C M L W Y K S Y N S N F S K T P W D I R

7561 GGGCCTGTCTGGTTGAAGACTGTAATGTATGTATGCTATGGTATAAAAGCTATAATTCAAATTTCAGTAAGACCCCTTGGGATATTAGAT

Y Q H L A M L K K M D V L M I H N A L V I T I K R K L L Y E

7651 ATCAACATCTAGCAATGTTAAAAAAAATGGATGTACTAATGATACACAATGCTCTTGTAATAACTATCAAAAGAAAATTGCTATACGAAT

**| ORF1 end**

* **| 3’LTR start**

7741 GATAGCTAAGAATCAAAAAGATAACAGGCAGGTTGTG**ATGGTGACGAGAGGGTG**TCGCAGGGCACAGAGCAGCCCTTGGTGGAGTAAGGA

**PPT**

7831 GCAATCTGACCACCGTCAAAGCCTCCCACGAACAAATAACAAATTGAGATAGGTTGCTAAACCACAAAGACAGACTCACATATGCTTGCT

7921 TCCTAAAGTAAGTTCAAATCCCCGAAACAGAAGATTTCCTGTTTGAACACCACTCGGTACATAAATATCAAACAATTTCTGTATTATGCT

8011 ACCTATGGAAAAGATGAAGGTGTATAAAAGTAAGATCGAGTGTAAAGCCAGCAGATCTCCAGCCCATTGGCACTTGGTTGCATCCAAGCA

8101 AGGAGACCTCTGGCTCAGTATGTAAATCTTATTGGTTAATGCTTGCTCAGGTCACTTAATATTTTATTATTCATTGTGTTTGATACTTCT

8191 GTATGACGTGTATATAGTAATAAATAGTAATTAATATATCTGCTCCTTGTGGCCTTGTAATTTATTGTCAGGGCCTTAATATAAGCAACC

8281 TGTGCGTAATCCCAACTATTCCTGATTGGCCCAGGCTAAGTAGTGGAATAAGTGATCTACGTGAGGCCTATACCCACGA**CA**

**3’LTR end |**

### ERV-Spuma.1-Cbo

1 **GT**AAATCTTATTATTTAATGCTTGCTCAGGTCACTTAACATTTTATTATTCATTGTGTTTGATACTTCTGTATGATGTGTATATGGTAAT

91 AAATAGTAATTAATATATCTGCTCCTTGTGGCCCTGTAATTTATTGTCAGGGGCTTAATGTAAGCAACTTGTGCGTAATCCCAACTATTC

181 CTGATTCGCCCAGGCCAAGTAATGGAATAAGTGATCTATGTGAGGCCTATACCCACAA**CA**AT**TGGCGCCCAATGTGGGC**TCCAGGAGCAC

**5′LTR end |** **PBS**

271 AATATATAACTGGTACCCTTAGAATAAATCCTAGATCTAGGCATGGCTCATAATCTGTATAATTTAGAGAGTTTTAATCAGAGGTTCAGA

**| Gag start**

M L Y Q R P P R H G E N I T I R I M N G P W R T G D R Y T Q

361 TGTTGTATCAACGCCCTCCAAGACACGGTGAAAATATCACCATACGCATTATGAATGGTCCATGGAGAACTGGTGATAGATATACACAAA

I T L E L Q D Q G G T N L P I P E Y A T L D G Q I E R T E I

451 TAACTTTAGAACTGCAAGATCAAGGAGGGACAAATTTGCCTATCCCCGAGTACGCAACACTGGACGGACAAATAGAAAGAACTGAAATTG

V I H A N F A E T R N W L G Q P P D I N T G V D R H G P M A

541 TAATTCATGCAAATTTTGCTGAAACGCGAAATTGGTTAGGCCAACCACCAGACATTAATACTGGAGTGGATCGACATGGACCAATGGCTC

H E P F P P R D E I C E G Y L P I T L E E L E Q L N Q P G K

631 ATGAACCTTTTCCTCCAAGAGATGAGATTTGTGAAGGATATTTACCTATAACCTTGGAAGAATTAGAGCAACTAAACCAACCAGGAAAAT

L R I E A A L L T R L Y S Q Q R T Q I G T N V G M G G A R I

721 TGAGAATAGAAGCAGCATTATTAACAAGATTATATAGTCAACAAAGGACACAAATAGGGACAAATGTTGGAATGGGAGGAGCTAGAATTC

P L V L V Q V A S L P F G N I R A A V G Q T P M D I K K I F

811 CATTAGTACTGGTCCAAGTTGCATCTCTCCCCTTTGGTAATATACGAGCAGCAGTGGGACAGACTCCTATGGATATTAAAAAGATATTTT

S W M A E R I N I L E G V L P N M N N A T R C Q L V N S L V

901 CCTGGATGGCTGAGAGGATTAATATATTAGAGGGGGTATTGCCCAATATGAATAATGCAACCAGATGTCAGCTGGTTAACTCACTAGTTC

P Y Q L S L N E E E C L S W D Q I I S C L Y A K A H R H I P

991 CCTATCAATTATCATTAAATGAAGAGGAATGTTTATCCTGGGACCAAATTATATCATGTTTATACGCTAAAGCTCATAGACATATTCCTA

T A K L G E E L Q R I S S E Q G Y K T A F S S G L A M T N Q

1081 CTGCTAAATTAGGAGAGGAATTACAGAGAATAAGCAGTGAACAAGGATACAAGACAGCATTCTCATCAGGATTAGCTATGACAAATCAGA

N Y G H L W G T V K N L V P G Q A P L A E I T C R L E V L P

1171 ATTATGGACATCTATGGGGAACTGTAAAAAATCTAGTTCCTGGTCAGGCTCCACTAGCAGAAATTACTTGCAGATTAGAAGTGTTACCAA

S D R E H I R Q F S T I V D T V Y R M L D L D P T G K M T G

1261 GTGACCGGGAACATATAAGGCAATTTTCAACTATAGTGGATACAGTCTACCGAATGTTAGACCTTGATCCTACAGGGAAAATGACAGGAA

```
     R  S  R  A  S  P  V  N  P  S  P  S  P  Q  G  K  S  R  N  K  K  P  F  K  K  R  F  P  K  E
1351 GATCAAGGGCTAGTCCAGTGAATCCCTCTCCCTCTCCTCAGGGCAAGAGTAGAAACAAGAAGCCTTTTAAGAAGAGATTTCCAAAAGAAA

     T  P  V  I  P  P  T  F  K  P  F  K  S  E  E  R  R  Y  P  A  S  Q  G  R  I  E  S  T  K  G
1441 CTCCAGTTATACCTCCAACATTCAAGCCTTTTAAATCAGAGGAGAGAAGGTATCCGGCATCTCAAGGAAGAATAGAATCTACTAAAGGAG

     A  Y  N  L  K  P  R  V  T  P  S  R  R  W  T  M  G  S  V  F  S  I  Q  L  P  W  A  F  C  P
1531 CTTATAACCTTAAACCCAGGGTTACTCCTTCACGACGATGGACAATGGGCTCAGTCTTCAGCATCCAATTGCCCTGGGCCTTCTGCCCCC
     Gag end |                                                                         Pol start |
     P  P  P  P  P  Q  G  *                                                                     M
1621 CCCCCCCCCCCCCCCCAGGGATAACAAGCCAGGTCCTTCGGGAATAAGGAAGAAAATTAACAGGATTAATAAAAATAGAAAGGACACATGA

     K  V  L  L  Q  I  Q  G  K  T  I  E  A  E  W  D  S  G  S  E  I  T  I  L  P  K  E  V  L  K
1711 AAGTTCTGCTCCAGATACAAGGAAAGACAATTGAGGCAGAATGGGACTCAGGATCAGAAATAACAATATTACCTAAAGAAGTGTTAAAAG

     G  L  L  P  I  K  K  N  K  L  M  T  I  T  G  E  T  E  V  P  V  F  Y  T  T  I  I  I  D  G
1801 GTCTCTTACCTATTAAAAAAAATAAACTAATGACCATTACTGGGGAGACTGAGGTACCTGTATTCTACACAACAATTATAATTGATGGGA

     K  N  R  R  I  E  V  V  E  S  P  D  G  Q  A  L  I  S  V  K  D  T  P  W  I  G  I  T  R  L
1891 AAAACAGAAGGATAGAAGTGGTTGAATCACCTGATGGACAAGCTTTAATTTCAGTGAAAGATACGCCTTGGATAGGTATAACCAGGCTAG
                                    Aspartic protease
     G  S  R  I  N  Y  Q  N  R  H  R  E  D  T  K  R  N  S  E  T  N  Q  F  V  R  S  R  G  K  A
1981 GAAGTAGAATTAACTATCAGAATCGACATCGAGAAGATACAAAGAGAAATTCTGAAACAAACCAATTTGTCAGATCGAGGGGAAAAGCAA

     I  G  T  I  F  L  S  F  C  P  L  V  A  E  M  G  K  S  G  R  T  Y  V  N  S  A  S  T  G  K
2071 TTGGAACAATTTTTTTATCATTTTGCCCCCTTGTGGCAGAAATGGGAAAATCAGGTAGGACATACGTCAATTCCGCCTCTACAGGGAAAA

     I  K  P  K  A  Q  K  Q  F  E  I  N  P  Q  A  I  P
                Q  K  Q  F  E  I  N  P  Q  A  I  P  S  I  Q  I  V  L  N  D  L  L  K  Q  S
2161 TAAAACCGAAAGCACAGAAACAGTTTGAAATAAATCCTCAAGCTATACCGTCTATACAAATAGTTCTAAATGACTTACTCAAACAAAGTG
                                 Reverse transcriptase domain
     V  L  R  Q  E  T  S  E  M  N  T  P  V  Y  P  V  P  K  G  E  G  K  R  R  L  V  L  D  Y  R
2251 TCCTCAGGCAAGAAACATCTGAAATGAATACACCAGTTTATCCAGTACCAAAAGGAGAGGGAAAGAGGAGATTAGTACTAGATTATAGAG

     A  M  N  K  V  T  P  A  I  A  A  Q  N  ----------------------------------------------------
2341 CCATGAATAAAGTAACCCCAGCCATAGCTGCACAAAATNNNNNNNNNNNNNNNNNNNNNNNNNNNNNNNNNNNNNNNNNNNNNNNNNNNN

     ------------------------------------------------------------------------------------------
2431 NNNNNNNNNNNNNNNNNNNNNNNNNNNNNNNNNNNNNNNNNNNNNNNNNNNNNNNNNNNNNNNNNNNNNNNNNNNNNNNNNNNNNNNNNN

     ------------------------------------------------------------------------------------------
2521 NNNNNNNNNNNNNNNNNNNNNNNNNTATTGCAAAGGAGCCGGATATATAGTTTCATTTCAAAAATCTGAAATAGGTAAGCATGAAGTAAACT

     F  L  G  F  V  I  T  N  E  G  R  G  L  T  D  Q  Y  K  E  K  L  L  N  L  Q  P  P  K  T  L
```

2611 TTTTGGGCTTTGTGATTACTAATGAGGGAAGAGGTCTAACAGATCAATATAAAGAAAAACTACTCAATTTACAGCCCCCAAAGACTCTTA

K Q L Q N I L G F L N F A C A F I S N F V E L V K P L H D A
2701 AGCAACTGCAGAATATTTTGGGATTTTTGAATTTTGCCTGTGCTTTTATTTCCAATTTTGTAGAATTGGTAAAACCTCTCCATGATGCCA

I I K A N N N E P F W E Q K Q Q D A L D G L I M A I N Q A A
2791 TCATTAAGGCAAATAACAATGAACCCTTTTGGGAACAAAAACAACAAGATGCTTTAGATGGATTAATCATGGCAATTAACCAAGCAGCTC

L L T E R D T T K P Q Q Q K T P F E L L F G L S K K V E F N
2881 TTTTAACTGAAAGAGACACAACCAAACCCCAGCAGCAAAAAACACCATTCGAACTACTCTTTGGGTTATCAAAGAAAGTAGAATTTAATC

L T S D L S R E E Q L A L L A E I R P M L A T T S T S S P L
2971 TTACTTCAGACCTCTCCAGAGAAGAACAATTAGCACTCCTAGCAGAAATACGACCTATGTTGGCGACTACCTCCACATCCTCGCCACTAC

**Pol end |**

P S P R S W R P L V G L L V Q E R V A T R R P L * P Q * K P
3061 CCTCACCTCGTTCATGGCGTCCTCTGGTTGGCCTTCTTGTCCAGGAGAGGGTAGCTACTCGACGACCATTATAACCTCAGTGAAAACCAC

3151 CAACTCCAATTATTAAGGTACTATCTGATCGAGTTGTTGAAATTGTGGATGAGAAGGGTAACCTGAGACAAGTATCAACAGATAATTTAA

3241 AAGTAACTCCACATCAGCAACAGAATGACCTTGTCATTGGACCAGTGGATGGCATGGGAGAGGGAAACTAGATGTAGAAAAGCACTTAAT

**| Env start**

M F Q N Y I E E L D E K I T W K T R C R
3331 ACCATCAGACAAGACAGAACATCCTAAAATAATGTTCCAGAATTATATTGAAGAATTGGATGAAAAAATAACATGGAAAACACGATGCAG

Y W G Y A A C A T S T R I A M Q I I F T L L I L M T M L G V
3421 ATATTGGGGCTATGCTGCATGTGCTACTAGTACTAGGATAGCAATGCAGATTATATTTACCTTACTCATATTAATGACTATGTTAGGAGT

T C T V I V R L Q W K Y A I E R Q G P T I T W N Q T I Y Q L
3511 AACTTGTACAGTAATAGTTAGGTTACAATGGAAATATGCTATAGAGAGACAAGGCCCTACAATTACTTGGAATCAAACAATATATCAATT

P S I H R I R R G L Y H H P L P V N V T I T G L K Q G L Y C
3601 ACCATCTATTCATAGAATTAGGAGAGGATTATATCATCATCCATTACCAGTAAATGTGACTATTACAGGATTAAAACAGGGACTATACTG

E P F P K L I V A K E R V L G I S Q I L I L D S D H M A E A
3691 CGAACCTTTTCCAAAACTGATAGTGGCCAAGGAGAGGGTGCTAGGCATATCCCAGATATTGATATTAGATTCGGATCATATGGCTGAAGC

N N L G D P V G K K I L I Q L L N E E M K Q L K D I T L S F
3781 AAACAATCTAGGAGATCCTGTTGGAAAAAAGATTCTTATTCAGCTCCTAAATGAGGAAATGAAACAGCTTAAAGATATTACTCTTAGTTT

E I P L D G P Q T Q Q E Y I Q K K C Y H E F A H C Y W I D Y
3871 TGAAATACCTTTAGATGGTCCACAAACTCAGCAAGAGTATATTCAAAAGAAATGTTACCATGAATTTGCACATTGTTACTGGATTGATTA

K E Q R K W P E P Q V I A D H C P H P R S G Y P R F S G K N
3961 TAAGGAGCAACGAAAATGGCCAGAACCCCAAGTAATAGCTGATCATTGCCCACACCCTAGAAGCGGATACCCTCGATTTTCTGGTAAAAA

Y W V E S P S S V T K Q E W E R L Q R L K Q G A R L Q T Y R

4051 TTACTGGGTAGAATCTCCTTCCTCTGTTACAAAACAGGAATGGGAAAGGCTACAAAGGTTAAAACAGGGAGCAAGGCTACAGACATATAG

L P G G H E T F T G A L M C T D D A Y H L W W S K N K T A Q

4141 GTTACCAGGGGGACATGAGACATTTACTGGAGCACTAATGTGTACAGATGATGCCTACCATTTATGGTGGAGCAAAAATAAAACAGCACA

K E L F L A Y V I Q I K K L I Q D T K T G K L K K V V L L N

4231 AAAAGAGCTATTCTTAGCCTATGTCATTCAAATAAAGAAGCTGATTCAAGATACAAAAACAGGGAAATTAAAAAAAGTTGTCCTTCTGAA

G W H D Q G K G K W F K S M D D L S F C R H P E L T V F L N

4321 TGGTTGGCATGATCAAGGAAAAGGGAAATGGTTCAAATCCATGGATGACTTGTCATTTTGTAGACATCCAGAACTTACAGTATTTCTGAA

G T Y Y K H S C T D G D C Q S T Q A N I T Q I K G C K N L T

4411 TGGCACATATTATAAACATTCATGCACGGACGGTGACTGTCAGAGCACTCAGGCAAATATAACACAGATAAAAGGTTGTAAAAACCTAAC

S S P K H P Y A C Q F C K A I Q N A T G E D F L Q L Q Y Y D

4501 AAGTTCACCTAAACACCCATATGCATGTCAATTTTGCAAGGCGATTCAAAATGCAACTGGGGAAGATTTTCTCCAACTGCAATATTATGA

Q Q Y L L Y P K Y S R M E E V S H T V D M G L L L H D H T F

4591 TCAGCAATATTTACTATACCCTAAATATAGCAGGATGGAAGAAGTCAGTCATACAGTAGATATGGGACTCCTTTTACATGATCACACATT

P G P W C V E A K Q V R Q Q N Y S L Y S L Y Q Q C L F H S Q

4681 TCCCGGACCCTGGTGTGTTGAAGCTAAGCAGGTCCGGCAACAGAATTATTCACTATATTCTTTATACCAGCAATGTTTATTTCACTCACA

K H P V D D V I S G M K Q R L F V Q K Q G E G Q P Y N L T T

4771 AAAACATCCAGTGGATGATGTAATTAGTGGAATGAAGCAAAGATTATTTGTACAAAAACAAGGAGAAGGGCAGCCCTACAACTTAACCAC

C Q P V S I L D I S R G R A I W G A N E T F L N Y T I Q T K

4861 GTGCCAGCCAGTCTCAATCTTAGACATATCACGAGGACGAGCTATCTGGGGCGCAAATGAAACCTTCCTAAATTATACAATACAGACTAA

K S K G C H T K Q K R S I T N L Q K I Q E A R L I L G S F I

4951 AAAATCAAAAGGCTGCCATACTAAGCAAAAGAGATCTATTACTAATCTACAAAAGATACAAGAGGCCAGATTGATCCTAGGTAGCTTCAT

T K I A K I S D L N D K E L S K G I H L L R D N L I T F A E

5041 TACTAAAATTGCCAAAATATCTGATTTAAATGATAAAGAATTATCAAAAGGGATACATCTGCTACGAGATAACCTTATTACATTTGCTGA

H T I D D V I Q M S Q T I T T V M I L S H I Q N L R I L L T

5131 ACATACAATTGATGATGTGATTCAAATGAGTCAAACCATTACCACAGTAATGATACTTTCTCATATACAGAATTTAAGAATCCTTTTAAC

E G K V D W N I L N S T W I Q E Q L R V T D E I I T L I R R

5221 TGAAGGCAAGGTTGATTGGAACATTTTAAACTCAACCTGGATACAAGAACAATTACGAGTCACAGATGAAATTATTACTTTAATAAGAAG

T A R G L A Y D I Q Q W V N K P Q K G V W E I S L Q Y D I V

5311 AACAGCTAGGGGATTAGCTTATGATATACAACAATGGGTTAACAAACCACAAAAAGGAGTCTGGGAAATCTCCCTACAGTATGACATTGT

I P Q R I Y P T N W K I I N Y G H L V Y T G N K G G R I W L

5401 TATACCACAAAGAATATATCCTACTAATTGGAAAATTATCAATTATGGTCATCTGGTATATACTGGAAATAAAGGAGGAAGAATCTGGTT

K H P Y T L I T Q G W G E V R Y L E V R Q C Y E Q D Y L I C

5491 AAAACATCCCTATACCCTAATAACCCAAGGGTGGGGGGAAGTAAGATATTTAGAAGTAAGACAATGCTATGAACAAGATTATCTTATTTG

D E V I E H E P C G N Q T G S R C P I M V E P I V S P Y L R

5581 TGATGAGGTCATTGAACATGAGCCATGTGGTAACCAAACTGGATCCAGATGTCCTATAATGGTAGAACCAATTGTCTCCCCATACCTTAG

I D L L K N G S Y I V M T S L E E C N V P P Y Q P S L I T V

5671 AATTGACCTATTGAAGAATGGGAGTTACATAGTGATGACCAGTTTGGAGGAGTGTAACGTACCACCGTACCAACCATCTCTGATCACAGT

H E T L T C Y G Y E F K P P L R K N T V I Q E N I P V P P L

5761 ACATGAGACACTGACTTGCTATGGATATGAATTTAAACCACCCCTTAGAAAAAACACAGTAATACAAGAAAATATACCTGTGCCACCACT

S V S L P H M I G I I A D L R R I K M E L A S S L G L S S R

5851 GTCAGTCAGCTTACCGCATATGATAGGTATTATAGCTGATCTCAGAAGAATAAAAATGGAGCTAGCTTCCTCTTTGGGACTCAGTTCACG

Y D Q E S K Y L T T E N R F T R R R R R I D V P Q R L N R L

5941 ATATGATCAAGAGAGCAAATACTTAACTACTGAGAATAGATTTACACGAAGGAGGAGAAGAATAGATGTACCTCAAAGGCTAAATAGACT

L E F I A D I W P A A A G A I K G I A N R I K D L T S G I F

6031 GTTAGAATTCATTGCAGACATCTGGCCTGCTGCAGCTGGAGCTATCAAGGGCATAGCTAATAGAATCAAGGATCTAACATCTGGAATTTT

G M A S D L L A Y A K P V I I G V V L L I I L V L V I K L I

6121 TGGTATGGCTTCTGATTTGTTAGCTTATGCAAAACCTGTGATAATAGGAGTGGTTTTGTTGATTATACTAGTGCTAGTCATCAAATTAAT

**Env end |**

S W I A T K C K T E *

6211 CAGTTGGATTGCTACAAAATGTAAAACAGAGTGAGATGATGGAATCACAAGATCAGGCTATGTTAGCAACGTTATGGTTTTACAGAGCTC

6301 TTACTGTAAAAAGTATACTTACTCATTATAAAACTATCCTGGTACCAATGCCGAGTTGTGACCAGGAGTCTACTTTTGTTTTTAGACACG

**Internal Promoter** **| ORF1 start**

M I P L P Q

6391 CTTGTCGTTATTGTGCTTATATCTATCATAGTAAATATCTAATTAAGAGAAGAGAAAATTGAGAAAGAGAATATGATACCACTACCCCAA

S H N S N I R D G I L P V H T L K P D T E I N H G Q Y P V I

6481 AGCCATAACTCAAACATACGGGATGGAATCCTACCTGTACACACATTAAAACCAGACACTGAAATCAATCATGGACAGTACCCTGTGATC

M Q L T L T A L E G A P P V W I Q T P A K E L A L I T L S D

6571 ATGCAACTGACCCTTACTGCTCTCGAAGGTGCACCACCTGTCTGGATACAGACTCCGGCGAAGGAACTAGCACTGATCACACTATCTGAC

R T K I L P S H M G W P I P M E Q P Q H L L Y F Q S L H H W

6661 AGGACTAAGATATTGCCCAGCCACATGGGATGGCCAATCCCAATGGAACAACCACAACATCTCCTATACTTTCAGAGTCTCCATCACTGG

Large insertion (CGATAACATATTCAAAGA)

M Y S S S S R N W M Q T S I T T H C L N L V F P P L G Q V Y

6751 ATGTACAGCAGCTCTTCGAGGAACTGGATGCAAACCTCGATCACCACTCATTGTCTGAATCTGGTATTCCCTCCCCTGGGGCAGGTTTAT

Insertion (A)

W F L T L T C C N I A S G L T H G H C Y M G L W E S G N N A

6841 TGGTTTTTAACGTTGACATGCTGTAACATAGCTAGTGGATTGACACATGGACATTGCTACATGGGTCTTTGGGAATCTGGAAACAATGCA

T A E G W L F W G R K P C L Q A Q G T S N L W C H K P L L L

6931 ACTGCTGAGGGATGGCTGTTCTGGGGTCGGAAACCATGCCTACAAGCACAAGGTACAAGTAATCTTTGGTGCCATAAGCCATTATTACTA

P M H C Y D T K F A F V S Y G A C L V E D C N L C M L W Y K

7021 CCCATGCATTGTTATGATACTAAATTTGCATTTGTTAGCTATGGGGCCTGTCTGGTTGAAGACTGTAATTTATGTATGCTATGGTATAAA

S Y N S N F S K T P W D I R Y Q H L A M L K K M D V L M I Y

7111 AGCTATAATTCAAATTTCAGTAAGACCCCTTGGGATATTAGATATCAACATCTAGCAATGTTAAAAAAAATGGATGTACTAATGATATAC

**ORF1 end |**

S A L V I T I K R K L L Y E *

7201 AGTGCTCTTGTAATAACTATCAAAAGAAAATTGCTATACGAATGATAGCTAAGAATCAAAAAGATAACAGCCAGGTTGTG**ATGGTGACGA**

**| 3′LTR start** **PPT**

7291 **GAGGGTG**TCGCAGGGCACAGAGCAGCCCTTGGTGGAGTAAGGAGCAATCTGACCACTGTCAAAGCCTCCCACGAACAAATAACAAATAGA

7381 GATAGGTTGCTAAACCACAAAGACAGACTCACATATGCTTGCTTCCTAATGTAAGTTCGAATCCCTGAAACAGAAGATTTCCTGTTTGAA

7471 CACCACTCGGTACATAAACATCAAACAATGTCTGTATTACGCTACCTATGGAAAAGATGAAGGTGTATAAAAGTAAGATCGAGTGTAAAG

7561 CCAGCAGATCTCCAGCCCATTGGCACTTGGTTGCATCCAAGCAAGGAGACCTCTGGCTCGGTATGTAAATCTTATTAGTTAATGCTTGCT

7651 CAGGTCACTTAATATTTTATTATTCATTGTGTTTGATAATTCTGTATGACGTGTATATAGTAATAAATAGTAATTAATATATCTGCTCCT

7741 TGTGGCCTTGTAATTTATTGTCAGGGCCTTAATGTAAGCAACCTGTGTGTAATCCCAACTATTCCTGATTGGCCCAGGCTAAGTAGTGGA

7831 ATAAGTGATCTACGTGAGGCCTATACCCACGA**CA**

**3′LTR end |**

**Fig. S1. Detailed descriptions of the putative genomes of ERV-Spuma.1-Cma, ERV-Spuma.2-Cma and ERV-Spuma.1-Cbo.** The location of the proteins encoded by the *gag*, *pol*, *env* and *ORF 1* genes were determined via homology to representative foamy viruses, searching CDD, and the distribution of start and stop codons, determined by ORFfinder (https://www.ncbi.nlm.nih.gov/orffinder/). The conserved domains were determined by searching against the CDD and Pfam databases and highlighted in darker colors. The bold nucleotide sequences on black lines indicate the primer binding site (PBS), internal promoter and poly-purine tract (PPT). The important insertions in ORF1 were highlighted. The inferred pre-substitution nucleotides are shown in bold red.

**Spuma virus Gag domain (pfam03276)**

**E-value: 8.99e-30**

```
ERV-Spuma.1-Cma   236 RPPTHGENITVCIMNGPWGIGDRYKRKRLELQDQGgaNLPIPQWQHQ---DGRIERTEIVIHANFAEtLNWLGQPPDINT 312
Cdd:pfam03276      20 RNPLHGEIIGLRLTEGWWGQLERFQMVRLILQDED--NEPLQRPRHEiipRAVNPHTMFVLSGPLAE-LQLAFQDLDLPE 96

                  313 GEDRHGPMAHEPFTPGDEICEGYLPITLEELEQ---------LNQPANLRIEAALLARLYS------QQRTQRRTNVGTG 377
                   97 GPLRFGPLANGHYVEGDPYSRSYRPVTMAETAQmtrdeledtLNTQSEIEIQMINLLELYEvetralRRQLAERSSIGQG 176

                  378 G------------------------------------------------------------------------------- 378
                  177 Gispgashsrppvssfsglpslpaipgihtrapsppratstpgniprslgddnmpsssfagpsqprvsfhpgnpfaeaeg 256

                  379 --------------ARAPLIPVQVTS-------------------LPFSNI*AAVGPTPMDIKNIFSWVAERINVLEGVL 425
                  257 hrprsqsrerrrdiPSAPVISAPVPSappmiqyipvpppppvgavIPIQHIRSVTGEPPRNPREIPIWLGRNAPAIDGVF 336

                  426 PHMNNAARRQVVNSLV--PYQLSLNEEECVSWD*MISCLYSKAHGHIPTAKLGEELQRISSEQGIKTAFQLGLAMTNQNY 503
                  337 PTTTPDLRCRIINALLggNLGLSLTPGDCITWDSAVATLFIRTYGQYPLHQLGNVLKGIADQEGVATAYTLGMMLSGQNY 416

                  504 GHVWRIIKNLVPGQAPLAEITRRLEALPSDQERIRQF*TIVDMVYRMLDLDPTGRrtgTSRASpVNPSPS-SQGKSKGKK 582
                  417 QLVSGIIRGYLPGQAVVTAMQQRLDQEIDDQTRAETFIQHLNAVYEILGLNARGQ---SIRAS-VTPQPRpSRGRGRGQS 492

                  583 pfmKVFPKEAPVIPPTFK----PFKPEERRYPASQGRTKPRKGGYNLRPRVTPPPQYG-----QWDQSLGSNRPGPSAPP 653
                  493 ---APEPSQGPVNSGRGRqcpaPGQNDRGSNIQNQGQENSSQGGYNLRSRTYQPQRYGggrgrRWNENTNNSETRPTEQS 569

                  654 PR 655
                  570 PQ 571
```

**Spuma aspartic protease (A9) domain (pfam03539)**

**E-value: 1.49e-04**

```
ERV-Spuma.1-Cma   739 ENRRI--ELAESPDGQALISAKDTPWIgvTRKEIELTIRIDIEKXQREILKQTDLSDQGKKQWEHILDRF 806
 Cdd:pfam03539     54 QGRKVeaEVIATPLDYVLIAPSDVPWY--KKKPLELTIKIDLEEQQETLLQQSALSKEGKELLKKLFLKY 121
```

(c)

**Foamy virus envelope protein domain (pfam03408)**

**E-value: 5.32e-39**

```
ERV-Spuma.1-Cma   1807 VSLDQGMAWERETRCRKAL-NLVPSDKTEQAKILFQNYNEESdeKITXATXCRYSGYAAYATSTRIAMWIIFTLLILMTM 1885
 Cdd:pfam03408       1 MTLQQWIIWNKMNKAHEALqNSTTVTDQQKEQIILEIQNEEV--RPTRKDKIRYLLYTCCATSSRVLAWMLLVCVLLIVV 78

                  1886 LGVTCTVIVRLQWKYAIERQGPTITWNQT---IHQSPSIHRIRRGL-YDHPL--LVNVPITGLKQGLY*EPFPKPIVAKE 1959
                    79 LVSCFLTISRIQWNRDIQVLGPVIDWNVTqraVYQPLQTRRIARSLrMQHPVpkYIEVNMTSIPQGVYYEPHPEPIVVTE 158

                  1960 RVLGISQILILDSDHMAEANNLGHPVgKEILTQLLNE---------IPFEIPLDGPQTQQEYMQKKRCHEFAHYYWIDYK 2030
                   159 RVLGLSQVLMINSENIANNANLTQEV-KKLLAEVVNEemqslsdvmIDFEIPLGDPRDQEQYIHRKCYQEFAHCYLVKYK 237

                  2031 EQ*KWPESQVIADHCP----HHGRGYPRFASEDYWVE 2063
                   238 TPKSWPTEGLIADQCPlpgyHAGLSYKPQSIWDYYIK 274
```

**E-value: 1.70e-10**

```
ERV-Spuma.1-Cma   2081 REQGYRHIGYxgdvrhllehsCMDDAYHSLWN-KNKMTQKELFLayvtqIKKLiQDMKTGK--LKKDALLNDWHDQGKGR 2157
Cdd:pfam03408      304 RQNNYSHVLF-----------CSDQLYSKWYNiENSIEQNEKFL-----LNKL-DNLTTGSslLKKRALPKEWSSQGKNA 366

                  2158 WFKPMDDLSFCRHPELTVFLNGTYHKHSCMEGDCQMTRANITHI 2201
                   367 LFKEINVLDVCSKPELVILLNTSYYSFSLWEGDCNFTKNMISQL 410
```

**E-value: 5.23e-111**

```
ERV-Spuma.1-Cma   2212 HKHPYACQFYRliQNATGEDFLQLXYYDQXYLLYPKYSRMEEVsrgVDTGLLLHDHTFPGFWCIEAKQVQXXNYSLYSLY 2291
Cdd:pfam03408      425 HMHPYACRFWR--SKNEKEETKCRPGEKEKCLYYPYQDSLEST---YDFGFLAYQKNFPAPICIEQQEIRDKDYEVYSLY 499

                  2292 QQCLFNSQKHPVDDVISGMKQRLSvqkqEEGYPCNlttcqpvsilDISGGQAIWGTNETFLNYTIVDTPKKPRGSYTKRK 2371
                   500 QECKLASKVHGIDTVLFSLKNFLN----HTGRPVN----------EMPNARAFVGLVDPKFPPSYPNVTREHYTSCNNRK 565

                  2372 RSTT--NLQKIQEAGLIPDSSITKTAKISDLNDK*LAKGLHLLRDHLITFPEHTIDVIQMSQSIIAVM-THSHIQNLRIL 2448
                   566 RRSTdnNYAKLKSMGYALTGAVQTLSQISDINDENLQQGIYLLRDHVITLMEATLHDISVMEGMFAVQhLHTHLNHLKTM 645

                  2449 LTEGKVDWDTLNSTWIQEQLRVSDEMMTLIRRTARGLAYDIQQRVDKPEKGVWEISLYYEIVIP*RIYSTNWKIINYGHL 2528
                   646 LLERRIDWTYMSSAWLQQQLQKSDDEMKVIKRIAKSLVYYVKQTYNSPTATAWEIGLYYELTIPKHVYLNNWNVVNIGHL 725

                  2529 VYTGNRGGRVWLKHPCTLITQGCGEVKYLEVRECYEQDYLICDEVIKHEPCGNQTG-SRCPIMVEPIVSPYLRIGPLKNG 2607
                   726 VQSAGQLTHVTIAHPYEIINKECTETKYLHLKDCRRQDYVICDVVEIVQPCGNSTDtSDCPVWAEAVKEPFVQVNPLKNG 805

                  2608 NYIVTTSLDECSIPPYQPSLITVNETVTCYGYEFKSPLRINTVIQENIHMPPLSVSLPHMIGIIADLRQIKIELASSWDS 2687
                   806 SYLVLASSTDCQIPPYVPSIVTVNETTSCYGLNFKKPLVAEERLGFEPRLPNLQLRLPHLVGIIAKIKGLKIEVTSSGES 885

                  2688 VHDVTERANTELLRIDLREGYTPQWLNRLSESIADIWP 2725
```

886 IKDQIERAKAELLRLDIHEGDTPAWIQQLAAATKDVWP 923

**Fig. S2. Conserved domain alignments of the original ERV-Spuma.1-Cma proteins and foamy virus.** The conserved domains were determined by searching the Conserved Domain Database (CDD). (a) The alignment of ERV-Spuma.1-Cma Gag proteins and Spuma virus Gag domain (pfam03276); (b) The alignment of ERV-Spuma.1-Cma pro proteins and Spuma aspartic protease (A9) domain (pfam03539); (c) The alignment of ERV-Spuma.1-Cma envelope proteins and foamy virus envelope protein domain (pfam03408). Numbers refer to the position in the ERV-Spuma.1-Cma each protein or conserved domain. Identical amino acid residues are highlighted in red, and grey and blue indicate gaps or different amino acid residues, respectively. The E-value was generated by Conserved Domain search.

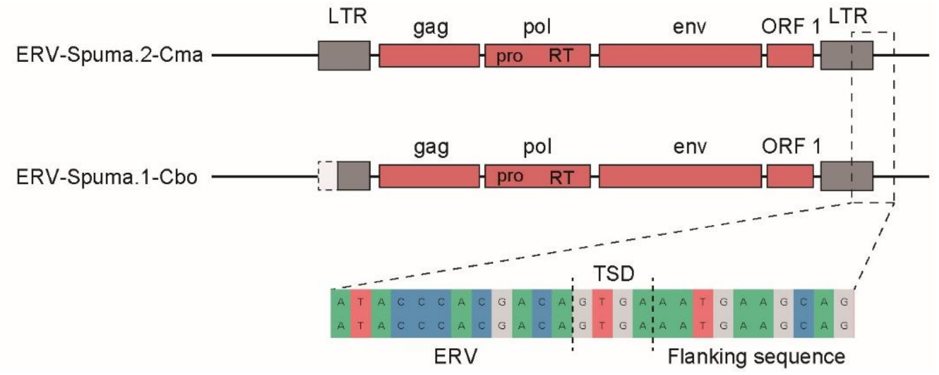


**Fig. S3. Comparison of two orthologous endogenous foamy viruses discovered in two bird species.** Dotted box indicates that 5’ LTR of ERV-Spuma.1-Cbo is defective. The alignment of 3’ LTR boundary of two EFVs is showed in detail and the alignment contains three parts, including 1) the end of the 3’LTR; 2) target site duplication; 3) flanking host sequences. Four base pair TSD directly following the intact bird EFV 5’ and 3’ LTR sequences. LTR, long terminal repeat; RT, reverse transcriptase; TSD, target site duplication.

**Table S1. Information on the 147 bird genomes in Genbank used for data mining.**

| No. | Species name | Common name | Accession no. |
|---|---|---|---|
| 1 | *Acanthisitta chloris* | Rifleman | GCA_000695815.1 |
| 2 | *Accipiter nisus* | Eurasian sparrowhawk | GCA_004320145.1 |
| 3 | *Acridotheres javanicus* | Javan myna | GCA_002849675.1 |
| 4 | *Agapornis roseicollis* | Peach-faced lovebird | GCA_002631895.1 |
| 5 | *Amazona aestiva* | Blue-fronted amazon | GCA_001420675.1 |
| 6 | *Amazona collaria* | Yellow-billed parrot | GCA_003947215.1 |
| 7 | *Amazona vittata* | Puerto Rican parrot | GCA_000332375.1 |
| 8 | *Anas platyrhynchos* | Mallard | GCA_003850225.1 |
| 9 | *Anas zonorhyncha* | Eastern spot-billed duck | GCA_002224875.1 |
| 10 | *Anser brachyrhynchus* | Pink-footed goose | GCA_002592135.1 |
| 11 | *Anser cygnoides domesticus* | Swan goose | GCA_000971095.1 |
| 12 | *Anser indicus* | Bar-headed goose | GCA_006229135.1 |
| 13 | *Antrostomus carolinensis* | Chuck-will's-widow | GCA_000700745.1 |
| 14 | *Apaloderma vittatum* | Bar-tailed trogon | GCA_000703405.1 |
| 15 | *Aptenodytes forsteri* | Emperor penguin | GCA_000699145.1 |
| 16 | *Apteryx australis mantelli* | Brown kiwi | GCA_001039765.1 |
| 17 | *Apteryx haastii* | Great spotted kiwi | GCA_003342985.1 |
| 18 | *Apteryx owenii* | Little spotted kiwi | GCA_003342965.1 |
| 19 | *Apteryx rowi* | Okarito brown kiwi | GCA_003343035.1 |
| 20 | *Aquila chrysaetos canadensis* | Golden eagle | GCA_000766835.1 |
| 21 | *Ara macao* | Scarlet macaw | GCA_000400695.1 |
| 22 | *Athene cunicularia* | Burrowing owl | GCA_003259725.1 |
| 23 | *Balearica regulorum gibbericeps* | East African grey crowned-crane | GCA_000709895.1 |
| 24 | *Bambusicola thoracicus* | Chinese bamboo-partridge | GCA_002909625.1 |
| 25 | *Bubo blakistoni* | Blakiston's fish-owl | GCA_004320225.1 |
| 26 | *Buceros rhinoceros silvestris* | Rhinoceros hornbill | GCA_000710305.1 |
| 27 | *Calidris pugnax* | Ruff | GCA_001431845.1 |
| 28 | *Calidris pygmaea* | Spoon-billed sandpiper | GCA_003697955.1 |
| 29 | *Callipepla squamata* | Scaled quail | GCA_002218305.1 |
| 30 | *Calypte anna* | Anna's hummingbird | GCA_000699085.1 |
| 31 | *Cariama cristata* | Red-legged seriema | GCA_000690535.1 |
| 32 | *Casuarius casuarius* | Southern cassowary | GCA_003342895.1 |
| 33 | *Cathartes aura* | Turkey vulture | GCA_000699945.1 |
| 34 | *Chaetura pelagica* | Chimney swift | GCA_000747805.1 |
| 35 | *Charadrius vociferus* | Killdeer | GCA_000708025.2 |
| 36 | *Chlamydotis macqueenii* | Macqueen's bustard | GCA_000695195.1 |
| 37 | *Chlamydotis undulata undulata* | Houbara bustard | GCA_003400225.1 |
| 38 | *Chrysolophus pictus* | Golden pheasant | GCA_003413605.1 |
| 39 | *Cicinnurus regius* | King bird of paradise | GCA_003713305.1 |
| 40 | *Ciconia boyciana* | Oriental stork | GCA_002002965.1 |
| 41 | *Colinus virginianus* | Northern bobwhite | GCA_000599465.2 |

| | | | |
|---|---|---|---|
| 42 | *Colius striatus* | Speckled mousebird | GCA_000690715.1 |
| 43 | *Columba livia* | Rock pigeon | GCA_000337935.1 |
| 44 | *Corapipo altera* | White-ruffed manakin | GCA_003945725.1 |
| 45 | *Corvus brachyrhynchos* | American crow | GCA_000691975.1 |
| 46 | *Corvus cornix cornix* | Hooded crow | GCA_000738735.2 |
| 47 | *Corvus hawaiiensis* | Hawaiian crow | GCA_003402825.1 |
| 48 | *Coturnix japonica* | Japanese quail | GCA_001577835.1 |
| 49 | *Crypturellus cinnamomeus* | Thicket tinamou | GCA_003342915.1 |
| 50 | *Cuculus canorus* | Common cuckoo | GCA_000709325.1 |
| 51 | *Cyanistes caeruleus* | Blue tit | GCA_002901205.1 |
| 52 | *Dendrocopos noguchii* | Okinawa woodpecker | GCA_004320165.1 |
| 53 | *Diphyllodes magnificus* | Magnificent bird-of-paradise | GCA_003713285.1 |
| 54 | *Dromaius novaehollandiae* | Emu | GCA_003342905.1 |
| 55 | *Egretta garzetta* | Little egret | GCA_000687185.1 |
| 56 | *Empidonax traillii* | Willow flycatcher | GCA_003031625.1 |
| 57 | *Eopsaltria australis* | Eastern yellow robin | GCA_003426825.1 |
| 58 | *Erythrura gouldiae* | Gouldian finch | GCA_003676055.1 |
| 59 | *Eudromia elegans* | Elegant crested-tinamou | GCA_003342815.1 |
| 60 | *Eurypyga helias* | Sunbittern | GCA_000690775.1 |
| 61 | *Falco cherrug* | Saker falcon | GCA_000337975.1 |
| 62 | *Falco peregrinus* | Peregrine falcon | GCA_000337955.1 |
| 63 | *Ficedula albicollis* | Collared flycatcher | GCA_000247815.2 |
| 64 | *Fulmarus glacialis* | Northern fulmar | GCA_000690835.1 |
| 65 | *Gallirallus okinawae* | Okinawa rail | GCA_002003005.1 |
| 66 | *Gallus gallus* | Chicken | GCA_000002315.5 |
| 67 | *Gavia stellata* | Red-throated loon | GCA_000690875.1 |
| 68 | *Geospiza fortis* | Medium ground-finch | GCA_000277835.1 |
| 69 | *Grus japonensis* | Red-crowned crane | GCA_002002985.1 |
| 70 | *Grus nigricollis* | Black-necked crane | GCA_004360235.1 |
| 71 | *Haliaeetus albicilla* | White-tailed eagle | GCA_000691405.1 |
| 72 | *Haliaeetus leucocephalus* | Bald eagle | GCA_000737465.1 |
| 73 | *Hemignathus virens* | Hawaii amakihi | GCA_003286495.1 |
| 74 | *Himantopus himantopus leucocephalus* | Black-winged stilt | GCA_003993805.1 |
| 75 | *Hirundo rustica rustica* | Barn swallow | GCA_003692655.1 |
| 76 | *Junco hyemalis* | Dark-eyed junco | GCA_003829775.1 |
| 77 | *Lagopus muta japonica* | Rock ptarmigan | GCA_004320205.1 |
| 78 | *Lepidothrix coronata* | Blue-crowned manakin | GCA_001604755.1 |
| 79 | *Leptosomus discolor* | Cuckoo roller | GCA_000691785.1 |
| 80 | *Limosa lapponica baueri* | Bar-tailed godwit | GCA_002844005.1 |
| 81 | *Lonchura striata domestica* | Bengalese finch | GCA_002197715.1 |
| 82 | *Lyrurus tetrix tetrix* | Black grouse | GCA_000586395.1 |
| 83 | *Manacus vitellinus* | Golden-collared manakin | GCA_001715985.2 |
| 84 | *Meleagris gallopavo* | Turkey | GCA_000146605.3 |

| | | | |
|---|---|---|---|
| 85 | *Melopsittacus undulatus* | Budgerigar | GCA_000238935.1 |
| 86 | *Merops nubicus* | Carmine bee-eater | GCA_000691845.1 |
| 87 | *Mesitornis unicolor* | Brown roatelo | GCA_000695765.1 |
| 88 | *Mixornis gularis* | Striped tit-babbler | GCA_003546035.1 |
| 89 | *Nannopterum auritus* | Double-crested cormorant | GCA_002173455.1 |
| 90 | *Nannopterum brasilianus* | Neotropic cormorant | GCA_002174335.1 |
| 91 | *Nannopterum harrisi* | Galapagos flightless cormorant | GCA_002173475.1 |
| 92 | *Neopelma chrysocephalum* | Saffron-crested tyrant-manakin | GCA_003984885.2 |
| 93 | *Nestor notabilis* | Kea | GCA_000696875.1 |
| 94 | *Nipponia nippon* | Crested ibis | GCA_000708225.1 |
| 95 | *Nothoprocta perdicaria* | Chilean tinamou | GCA_003342845.1 |
| 96 | *Numida meleagris* | Helmeted guineafowl | GCA_002078875.2 |
| 97 | *Opisthocomus hoazin* | Hoatzin | GCA_000692075.1 |
| 98 | *Paradisaea raggiana* | Raggiana bird of paradise | GCA_003713265.1 |
| 99 | *Paradisaea rubra* | Red bird of paradise | GCA_003713215.1 |
| 100 | *Parotia lawesii* | Lawes's parotia | GCA_003713295.1 |
| 101 | *Parus major* | Great tit | GCA_001522545.2 |
| 102 | *Passer domesticus* | House sparrow | GCA_001700915.1 |
| 103 | *Patagioenas fasciata monilis* | Band-tailed pigeon | GCA_002029285.1 |
| 104 | *Pavo cristatus* | Indian peafowl | GCA_005519975.1 |
| 105 | *Pelecanus crispus* | Dalmatian pelican | GCA_000687375.1 |
| 106 | *Phaethon lepturus* | White-tailed tropicbird | GCA_000687285.1 |
| 107 | *Phalacrocorax carbo* | Great cormorant | GCA_000708925.1 |
| 108 | *Phasianus colchicus* | Ring-necked pheasant | GCA_004143745.1 |
| 109 | *Phoenicopterus ruber ruber* | American flamingo | GCA_000687265.1 |
| 110 | *Phylloscopus plumbeitarsus* | Two-barred warbler | GCA_001655115.1 |
| 111 | *Phylloscopus trochiloides viridanus* | Greenish warbler | GCA_001655095.1 |
| 112 | *Phylloscopus trochilus acredula* | Willow warbler | GCA_002305835.1 |
| 113 | *Picoides pubescens* | Downy woodpecker | GCA_000699005.1 |
| 114 | *Pipra filicauda* | Wire-tailed manakin | GCA_003945595.1 |
| 115 | *Podiceps cristatus* | Great crested grebe | GCA_000699545.1 |
| 116 | *Pseudopodoces humilis* | Tibetan ground-tit | GCA_000331425.1 |
| 117 | *Psittacula krameri* | Rose-ringed parakeet | GCA_002870145.1 |
| 118 | *Pterocles gutturalis* | Yellow-throated sandgrouse | GCA_000699245.1 |
| 119 | *Pterocnemia pennata* | Lesser rhea | GCA_003342835.1 |
| 120 | *Pygoscelis adeliae* | Adelie penguin | GCA_000699105.1 |
| 121 | *Pygoscelis antarcticus* | Chinstrap penguin | GCA_003264595.1 |
| 122 | *Pygoscelis papua* | Gentoo penguin | GCA_003264615.1 |
| 123 | *Recurvirostra avosetta* | Pied Avocet | GCA_004023745.1 |
| 124 | *Rhea americana* | Greater rhea | GCA_003343005.1 |
| 125 | *Saxicola maurus maurus* | Siberian stonechat | GCA_900205225.1 |
| 126 | *Scolopax mira* | Amami woodcock | GCA_004320125.1 |
| 127 | *Serinus canaria* | Common canary | GCA_000534875.1 |
| 128 | *Setophaga coronata coronata* | Yellow-rumped warbler | GCA_001746935.1 |

| | | | |
|---|---|---|---|
| 129 | *Spheniscus humboldti* | Humboldt's penguin | GCA_003264545.1 |
| 130 | *Spheniscus magellanicus* | Magellanic penguin | GCA_003264715.1 |
| 131 | *Spheniscus mendiculus* | Galapagos penguin | GCA_003264655.1 |
| 132 | *Sporophila hypoxantha* | Tawny-bellied seedeater | GCA_002167245.1 |
| 133 | *Streptopelia turtur* | European Turtle-dove | GCA_901699155.1 |
| 134 | *Strigops habroptila* | Kakapo | GCA_004027225.1 |
| 135 | *Strix occidentalis caurina* | Spotted owl | GCA_002372975.1 |
| 136 | *Struthio camelus australis* | African ostrich | GCA_000698965.1 |
| 137 | *Sturnus vulgaris* | Common starling | GCA_001447265.1 |
| 138 | *Syrmaticus mikado* | Mikado pheasant | GCA_003435085.1 |
| 139 | *Taeniopygia guttata* | Zebra finch | GCA_000151805.2 |
| 140 | *Tauraco erythrolophus* | Red-crested turaco | GCA_000709365.1 |
| 141 | *Tinamus guttatus* | White-throated tinamou | GCA_000705375.2 |
| 142 | *Tympanuchus cupido pinnatus* | Greater prairie chicken | GCA_001870855.1 |
| 143 | *Tyto alba* | Barn owl | GCA_000687205.1 |
| 144 | *Uria lomvia* | Thick-billed guillemot | GCA_002289315.1 |
| 145 | *Urile pelagicus* | Pelagic cormorant | GCA_002173435.1 |
| 146 | *Zonotrichia albicollis* | White-throated sparrow | GCA_000385455.1 |
| 147 | *Zosterops lateralis melanops* | Silver-eye | GCA_001281735.1 |

**Table S2. Information on the representative retroviruses.**

| No. | Virus name | Genus | Abbreviation | Natural host | Accession no. |
|---|---|---|---|---|---|
| 1 | Avian leukemia virus | Alpharetrovirus | ALV | Chicken | NC_015116 |
| 2 | Lymphoproliferative disease virus | Alpharetrovirus | LDV | Turkey | U09568 |
| 3 | Mouse mammary tumor virus | Betaretrovirus | MMTV | Mouse | NC_001503 |
| 4 | Mason-Pfizer monkey virus | Betaretrovirus | MPMV | Primate | NC_001550 |
| 5 | Simian retrovirus 1 | Betaretrovirus | SRV1 | Primate | M11841 |
| 6 | Bovine leukemia virus | Deltaretrovirus | BLV | Cattle | NC_001414 |
| 7 | Human T-lymphotropic virus 1 | Deltaretrovirus | HTLV1 | Human | NC_001436 |
| 8 | Simian T-lymphotropic virus 2 | Deltaretrovirus | STLV2 | Non-human primate | NC_001815 |
| 9 | Walleye dermal sarcoma virus | Epsilonretrovirus | WDSV | Fish | NC_001867 |
| 10 | Walleye epidermal hyperplasia virus type 1 | Epsilonretrovirus | WEHV1 | Fish | AF133051 |
| 11 | Walleye epidermal hyperplasia virus type 2 | Epsilonretrovirus | WEHV2 | Fish | AF133052 |
| 12 | Atlantic salmon swim bladder sarcoma virus | Gamma-epsilon | SSSV | Atlantic salmon | NC_007654 |
| 13 | Feline leukemia virus | Gammaretrovirus | FeLV | Cat | NC_001940 |
| 14 | Friend murine leukemia virus | Gammaretrovirus | F-MuLV | Mouse | NC_001362 |
| 15 | Mus dunni endogenous retrovirus | Gammaretrovirus | MDEV | Mouse | AF053745 |
| 16 | Porcine endogenous retrovirus A | Gammaretrovirus | PERV-A | Pig | AJ293656 |
| 17 | Rhinolophus ferrumequinum retrovirus | Gammaretrovirus | RfRV | Greater horseshoe bat | JQ303225 |
| 18 | Equine infectious anemia virus | Lentivirus | EIAV | Horse | NC_001450 |
| 19 | Feline immunodeficiency virus | Lentivirus | FIV | Cat | NC_001482 |
| 20 | Human immunodeficiency virus 1 | Lentivirus | HIV1 | Human | NC_001802 |
| 21 | Visna/Maedi virus | Lentivirus | VMV | Sheep | NC_001452 |
| 22 | Bovine foamy virus | Spumavirus | BFVbta | Cattle | NC_001831 |
| 23 | Equine foamy virus | Spumavirus | EFVeca | Horse | NC_002201 |
| 24 | Feline foamy virus | Spumavirus | FFVfca | Cat | NC_001871 |
| 25 | Brown greater galago prosimian foamy virus | Spumavirus | SFVocr | Greater galago | KM233624 |
| 26 | White-tufted-ear marmoset simian foamy virus | Spumavirus | SFVcja | Common marmoset | GU356395 |
| 27 | Squirrel monkey simian foamy virus | Spumavirus | SFVssc | Squirrel monkey | GU356394 |
| 28 | Orangutan Simian foamy virus | Spumavirus | SFVppy | Pongo pygmaeus pygmaeus | AJ544579 |

| | | | | | |
|---|---|---|---|---|---|
| 29 | Macaque simian foamy virus | Spumavirus | SFVmcy | Macaque | NC_010819 |
| 30 | African green monkey simian foamy virus | Spumavirus | SFVcae | African green monkey | NC_010820 |
| 31 | Western chimpanzee simian foamy virus | Spumavirus | SFVpve | Western chimpanzee | NC_001364 |
| 32 | Western lowland gorilla simian foamy virus | Spumavirus | SFVggo | Western lowland gorilla | NC_039029 |
| 33 | Spider monkey simian foamy virus | Spumavirus | SFVaxx | Spider monkey | NC_039027 |
| 34 | Sloth endogenous foamy virus | Spumavirus | SloEFV | Sloth | ABVD02350954 |
| 35 | Sphenodon punctatus endogenous foamy virus | Spumavirus | ERV-Spuma-Spu | Tuatara | Ref. 1 |
| 36 | Coelacanth endogenous foamy-like virus | Spumavirus | CoeEFV | Coelacanth | Ref. 2 |
| 37 | Amphilophus citrinellus fomy-like virus | Spumavirus | AciFLERV_1 | Midas cichlid | CCOE01002251 |
| 38 | Amphilophus citrinellus fomy-like virus | Spumavirus | AciFLERV_2 | Midas cichlid | CCOE01002087 |
| 39 | Austrofundulus limnaeus fomy-like virus | Spumavirus | AliFLERV | Annual killifish | Ref. 3 |
| 40 | Notophthalmus viridescens | Spumavirus | NviFLERV | Eastern newt | Ref. 3 |
| 41 | latyfish endogenous retrovirus | Spumavirus | PlatyfishEFV | Platyfish | Ref. 4 |
| 42 | Danio rerio foamy virus | Spumavirus | DrFV-1 | Zebrafish | CABZ01054182 |
| 43 | Cynops pyrrhogaster fomy-like virus | Spumavirus | CpyFLERV_1 | Japanese fire belly newt | FS313726 |
| 44 | Cynops pyrrhogaster fomy-like virus | Spumavirus | CpyFLERV_2 | Japanese fire belly newt | FS296312 |
| 45 | Pleurodeles waltl fomy-like virus | Spumavirus | PwaFLERV | Iberian ribbed newt | JG015238 |
| 46 | Poecilia reticulata fomy-like virus | Spumavirus | PreFLERV | Guppy | AZHG01028727 |
| 47 | Poecilia formosa fomy-like virus | Spumavirus | PfoFLERV_1 | Amazon molly | AYCK01023761 |
| 48 | Poecilia formosa fomy-like virus | Spumavirus | PfoFLERV_2 | Amazon molly | AYCK01027102 |
| 49 | Larimichthys crocea fomy-like virus | Spumavirus | LcrFLERV | Large yellow croaker | JRPU01021077 |
| 50 | Stegastes partitus fomy-like virus | Spumavirus | SpaFLERV | Bicolor damselfish | JMKM01038484 |
| 51 | Fundulus heteroclitus fomy-like virus | Spumavirus | FheFLERV | Mummichog | JXMV01100753 |
| 52 | Lates calcarifer fomy-like virus | Spumavirus | LcaFLERV | Barramundi | LBLR01010097 |
| 53 | Gadus morhua fomy-like virus | Spumavirus | GmoFLERV_1 | Atlantic cod | CAEA01131311 |
| 54 | Gadus morhua fomy-like virus | Spumavirus | GmoFLERV_2 | Atlantic cod | CAEA01539013 |
| 55 | Dicentrarchus labrax fomy-like virus | Spumavirus | DlaFLERV | European bass | CBXY010016181 |
| 56 | Oreochromis niloticus fomy-like virus | Spumavirus | OniFLERV | Nile tilapia | AERX01018483 |
| 57 | Cynoglossus semilaevis fomy-like virus | Spumavirus | CseFLERV | Tongue sole | AGRG01002780 |
| 58 | Sebastes rubrivinctus fomy-like virus | Spumavirus | SruFLERV | Flag rockfish | AUPQ01030678 |
| 59 | Sebastes nigrocinctus fomy-like virus | Spumavirus | SniFLERV | Tiger rockfish | AUPR01019601 |

| | | | | | |
|---|---|---|---|---|---|
| 60 | Pimephales promelas fomy-like virus | Spumavirus | PprFLERV_1 | Fathead minnow | JNCD01073002 |
| 61 | Pimephales promelas fomy-like virus | Spumavirus | PprFLERV_2 | Fathead minnow | JNCD01029789 |
| 62 | Thunnus orientalis fomy-like virus | Spumavirus | TorFLERV | Pacific bluefin tuna | BADN01112239 |
| 63 | Periophthalmus magnuspinnatus fomy-like virus | Spumavirus | PmaFLERV | Mudskipper | JACL01052273 |
| 64 | Periophthalmodon schlosseri fomy-like virus | Spumavirus | PscFLERV | Mudskipper | JACM01000693 |
| 65 | Anoplopoma fimbria fomy-like virus | Spumavirus | AfiFLERV | Sablefish | AWGY01041462 |
| 66 | Cyprinus carpio fomy-like virus | Spumavirus | CcaFLERV | Common carp | LN590673 |
| 67 | Nothobranchius furzeri fomy-like virus | Spumavirus | NfuFLERV | Turquoise killifish | JNBZ01063262 |
| 68 | Esox lucius fomy-like virus | Spumavirus | EluFLERV | Common pike | AZJR02000232 |
| 69 | Callorhinchusmilii fomy-like virus | Spumavirus | CmiFLERV_1 | Australian ghostshark | XM_007890932 |
| 70 | Callorhinchusmilii fomy-like virus | Spumavirus | CmiFLERV_2 | Australian ghostshark | AAVX02030290 |
| 71 | Hynobius retardatus fomy-like virus | Spumavirus | HreFLERV | Hokkaido salamander | LE148029 |
| 72 | Snakehead retrovirus | Unclassified | SnRV | Fish (snakehead fish) | NC_001724 |

**Table S3. The significant hits identified in *Ciconia maguari* and *Ciconia boyciana* genomes**

| Species name | Contig name | Contig size (bp) | Genomic region present | Top Blastp hit | Blastp E-value |
|---|---|---|---|---|---|
| *Ciconia maguari* | scaffold1426 | 348,633 | gag | gag [Feline foamy virus]_YP_009513248.1 | 2E-42 |
| | | | pol | pol [Rhesus macaque simian foamy virus]_YP_009513242.1 | 0.0 |
| | | | env | Env [Equine foamy virus]_ NP_054717.1 | 3E-139 |
| | scaffold3233 | 119,292 | gag | gag polyprotein [Feline foamy virus]_AAC58530.1 | 6E-39 |
| | | | pol | Pol [Simian foamy virus 3]_ATW64776.1 | 5E-131 |
| | | | env | Env [Equine foamy virus]_ NP_054717.1 | 0.0 |
| | scaffold2698 | 251,554 | pol | pol protein [Yellow-breasted capuchin simian foamy virus]_YP_009508582.1 | 8E-179 |
| | | | env | Env [Equine foamy virus]_ NP_054717.1 | 1E-136 |
| | scaffold291 | 44,351 | gag | gag polyprotein [Feline foamy virus]_AAC58530.1 | 1E-36 |
| | scaffold17755 | 1,078 | pol | pol protein [Human spumaretrovirus]_AAA66556.1 | 1E-59 |
| | scaffold15300 | 513 | env | Env [Equine foamy virus]_ NP_054717.1 | 6E-53 |
| | scaffold20855 | 498 | pol | pol protein [Yellow-breasted capuchin simian foamy virus]_YP_009508582.1 | 2E-35 |
| | C9153445 52.0 | 250 | pol | Pol [Equine foamy virus]_NP_054716.1 | 1E-30 |
| | C8901342 | 149 | env | env protein [Guenon simian foamy virus]_YP_009666127.1 | 9E-06 |
| | C8834376 | 134 | env | envelope protein, partial [Human spumaretrovirus]_AAF00493.1 | 2E-05 |
| | C8812898 | 130 | env | env [Spider monkey simian foamy virus]_YP_009508562.1 | 3E-02 |
| *Ciconia boyciana* | BDFF02011124 | 75,962 | gag | Gag [Puma feline foamy virus]_YP_009508536.1 | 1E-41 |
| | | | pol | Pol [Simian foamy virus 3]_ATW64776.1 | 1e-39 |
| | | | env | envelope protein [Bovine foamy virus]_AWK77175.1 | 0.0 |
| | BDFF02022209 | 7,476 | gag | gag [Feline foamy virus]_YP_009513248.1 | 3E-42 |
| | BDFF02022208 | 3,643 | pol | pol protein [Yellow-breasted capuchin simian foamy virus]_YP_009508582.1 | 0.0 |
| | | | env | Env [Equine foamy virus]_ NP_054717.1 | 2E-22 |

| | | | | | |
|---|---|---|---|---|---|
| | BDFF02034589 | 3,508 | pol | pol protein [Yellow-breasted capuchin simian foamy virus]_YP_009508582.1 | 0.0 |
| | | | env | envelope protein [Bovine foamy virus]_AWK77175.1 | 3E-136 |
| | BDFF02030469 | 2,952 | gag | Gag [Puma feline foamy virus]_YP_009508536.1 | 1E-31 |
| | | | pol | pol [Feline foamy virus]_YP_009513249.1 | 6E-82 |
| | BDFF02044953 | 728 | env | Env [Equine foamy virus]_ NP_054717.1 | 8E-71 |
| | BDFF02043489 | 728 | env | Env [Equine foamy virus]_ NP_054717.1 | 6E-69 |

**Table S4. The endogenous foamy viruses identified in *Ciconia maguari* and *Ciconia boyciana* genomes.**

| Species name | ERV name | Contig number | Contig size (bp) | Location (start-end) | Genomic region present |
|---|---|---|---|---|---|
| *Ciconia maguari* | ERV-Spuma.1-Cma | scaffold1426 | 348,633 | 684-10668 | 5'LTR-gag-pol-env-3'LTR |
| *Ciconia maguari* | ERV-Spuma.2-Cma | scaffold3233 | 119,292 | 59220-67580 | 5'LTR-gag-pol-env-3'LTR |
| *Ciconia maguari* | ERV-Spuma.3-Cma | scaffold2698 | 251,554 | 1-6733 | pol-env-3'LTR |
| *Ciconia maguari* | ERV-Spuma.4-Cma | scaffold291 | 44,351 | 42319-44351 | 5'LTR-Gag |
| *Ciconia boyciana* | ERV-Spuma.1-Cbo | BDFF02011124 | 75,962 | 16052-23912 | 5'LTR-gag-pol-env-3'LTR |

**Table S5. Dating the ERVs-Spuma-Cma insertion based on LTR-LTR divergence.**

| Contig number | Divergence | Integration time (MYA) |
|---|---|---|
| ERV-Spuma.1-Cma | 0.012 | 3.16 |
| ERV-Spuma.2-Cma | 0.053 | 13.95 |
